# Correspondence of fentanyl brain pharmacokinetics and behavior measured via engineering opioids biosensors and computational ethology

**DOI:** 10.1101/2024.03.15.584894

**Authors:** Anand K. Muthusamy, Matthew H. Rosenberg, Charlene H. Kim, Alex Z. Wang, Haruka Ebisu, Theodore M. Chin, Ashil Koranne, Jonathan S. Marvin, Bruce N. Cohen, Loren L. Looger, Yuki Oka, Markus Meister, Henry A. Lester

**Affiliations:** Division of Biology and Biological Engineering, California Institute of Technology; Division of Chemistry and Chemical Engineering, California Institute of Technology; Janelia Research Campus, Howard Hughes Medical Institute; University of California San Diego

## Abstract

Despite the ongoing epidemic of opioid use disorder and death by fentanyl overdose, opioids remain the gold standard for analgesics. Pharmacokinetics (PK) dictates the individual’s experience and utility of drugs; however, PK and behavioral outcomes have been conventionally studied in separate groups, even in preclinical models. To bridge this gap, we developed the first class of sensitive, selective, and genetically encodable fluorescent opioid biosensors, iOpioidSnFRs, including the fentanyl sensor, iFentanylSnFR. We expressed iFentanylSnFR in the ventral tegmental area of mice and recorded [fentanyl] alongside videos of behaviors before and after administration. We developed a machine vision routine to quantify the effects of the behavior on locomotor activity. We found that mice receiving fentanyl exhibited a repetitive locomotor pattern that paralleled the [fentanyl] time course. In a separate experiment, mice navigating a complex maze for water showed a dose-dependent impairment in navigation, in which animals repeated incorrect paths to the exclusion of most of the unexplored maze for the duration of the average fentanyl time course. This approach complements classical operant conditioning experiments and introduces a key feature of human addiction, the ability to carry out an ethologically relevant survival task, only now quantified in rodents. Finally, we demonstrate the utility of iFentanylSnFR in detecting fentanyl spiked into human biofluids and the generalizability of engineering methods to evolve selective biosensors of other opioids, such as tapentadol and levorphanol. These results encourage diagnostic and continuous monitoring approaches to personalizing opioid regimens for humans.

## Introduction

Humans have used opioids from plants for over 5,000 years and continue to consume them for their euphoric and analgesic properties^1^. Since ∼1870, the development of even more potent opioid agonists, such as heroin and fentanyl, has driven an increase in opioid use disorder (OUD). Today, OUD affects ∼16 million people worldwide, including ∼2 million in the U.S.^2^. At least 50% of these cases begin with an opioid prescription^3–6^. Among the ∼110,000 drug overdose deaths in the United States in the past year, fentanyl and other synthetic opioids alone account for ∼70%^7^.

Still, opioids remain the gold standard for severe and chronic pain despite the burden of OUD. Conventional opioids provide profound and general analgesic effects useful in severe pain, chronic pain (e.g., from cancer), some degenerative conditions, and palliative care. These opioids are unmatched by other classes of analgesics in not only blunting not only physical pain but also the perceptual and emotional factors of pain^8–10^. A core tenet of behavioral neuropharmacology is the existence of some mapping between the time course of a drug and the behavioral outcomes. Interindividual variability in pharmacokinetics, partly driven by genetic variance in metabolic enzymes, complicates the problem of optimized opioid dosing, especially outside the clinic^11^. The problem of personalizing pharmacokinetics is severe in substance abuse disorders: the patient must receive levels that relieve pain, minimize tolerance and other side effects, and remain within a therapeutic window to maximize adherence. The ideal window is a “moving target” tolerance that leads to a decreased response to the drug.

A central tension in opioid pharmacology lies in the differing tolerance rates for the *µ*-opioid receptor’s activity in circuits driving reward, analgesia, and respiratory depression^12^. These differences lead to withdrawal periods that decrease quality of life and dose escalation that can lead to undesirable side effects and, potentially, death by overdose. The mechanisms of tolerance and dependence are incompletely understood, but cellular neurobiology reveals a “location bias”: the cellular compartment in which a drug interacts with the *µ*-opioid receptor affects signaling dynamics^13–15^. Circuit adaptations are sensitive to variations in opioid dose regimens and their interruptions^16–18^. Therefore, relevant progress in neurobiology requires improved methods of monitoring opioids and their receptors across time and space^19–21^. Along the range from organelle to behaving animals, one asks, “What is the time course of the drug in the relevant compartment?” followed by “How does the cell or animal respond?”. In this work, we developed iOpioidSnFRs for sufficient sensitivity, selectivity, and kinetics to enable pharmacokinetic measurements at both subcellular and whole animal levels (**Figure 1A, B**).

**Figure 1:**
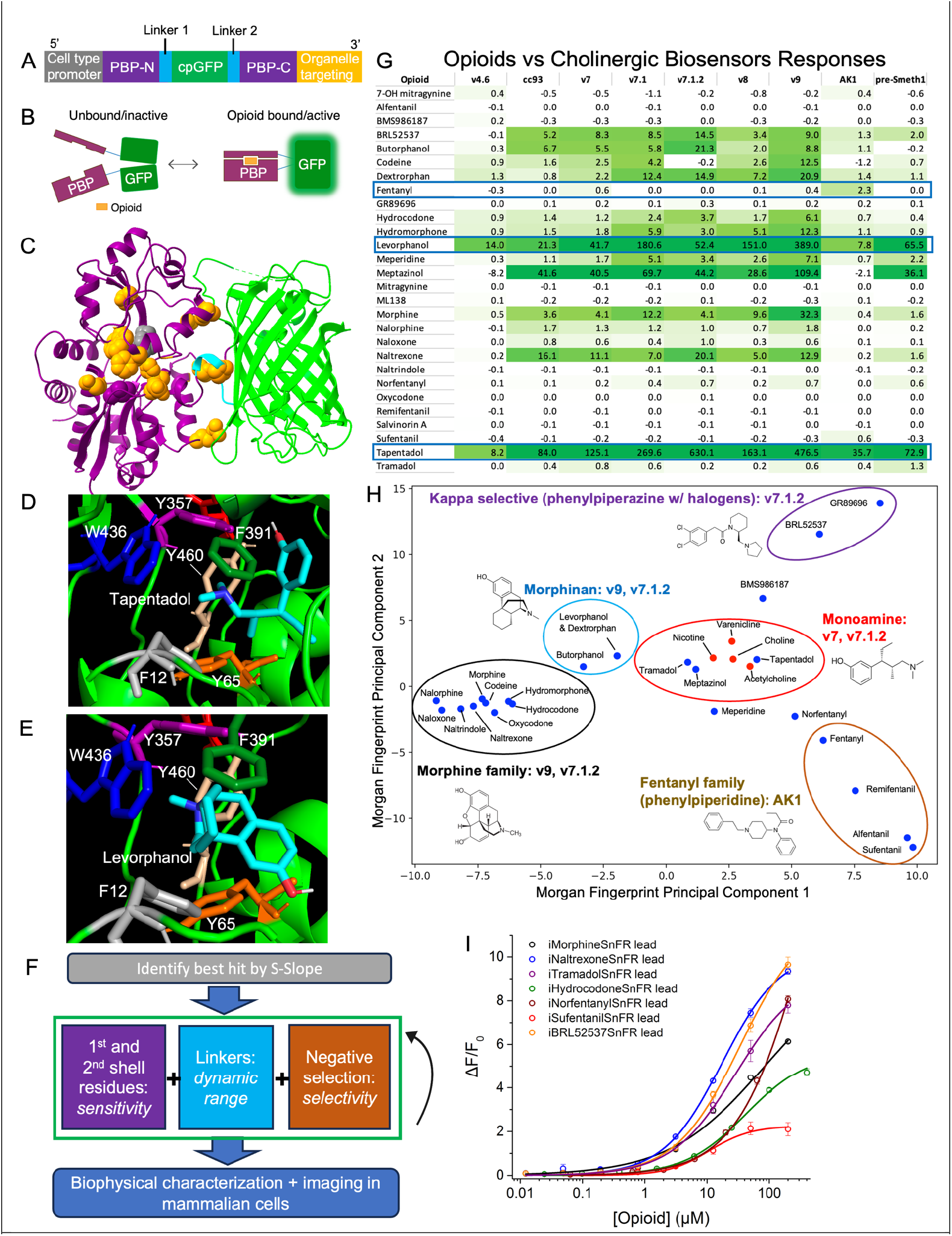
A General Strategy to Evolve Opioid Biosensors from Cholinergic Biosensors. (A) The biosensor gene construct. The periplasmic binding protein (PBP) gene (purple) is interrupted by a circularly permuted GFP (green), connected by two linker sequences (cyan). A 5’ promoter sequence restricts expression to the desired cell type, and a C-terminal tag directs the biosensor to the desired cellular compartment. (B) Scheme for ligand gating of GFP fluorescence. (C) Crystal structure of iNicSnFR3a (PDB: 7S7V) annotated with variable regions exploited in directed evolution: sites mutated to improve cholinergic binding (orange spheres) and a critical cation-π residue, Y357 (grey spheres). (D & E) Docking of two exemplar opioids, tapentadol, and levorphanol, in iNicSnFR3a (PDB: 7S7V) using AutoDock Vina. Binding pocket residues (labeled) form an aromatic box. (F) General strategy for the directed evolution of iOpioidSnFRs (G) “Green map” map of the response of cholinergic biosensors to opioids expressed as the S-Slope x 100 (0-4, white to light green, and 4-40+, light green to dark green) (n = 3 dose responses averaged). (H) Mapping the OpuBC mutant hit sequences to opioid structural classes (encircled and given representative structures). Opioids (blue) and cholinergic ligands (red) were represented as Morgan fingerprint vectors and subjected to a principal component analysis. Mitragynine and 7-OH mitragynine were omitted for their outlying alkaloid structures, atypical of clinically used opioids. (I) After one to three rounds of directed evolution, iOpioidSnFR leads display detection limits of < ∼2 *µ*M in bacterial lysates. SEM is shown as error bars (n = 3 dose responses).

Rodent models are critical for therapeutic development, conserving several receptors and circuits involved in biological phenomena related to OUD^22^. Established paradigms show that *µ*-opioids elicit analgesic, rewarding, reinforcing, place preferent, anxiolytic, respiratory depressive, and hyperactive effects in rodents^23,24^. The diversity of behavioral profiles and susceptibility to developing OUD await explanation in both human and animal models^25,26^. Still, there is a need for research in the endophenotypes beyond self-administration for each model organism. The field has established the positive valence of opioid effects; one now asks how the drugs affect higher-order behaviors that lead to and constitute OUD^27^.

The next critical open question is when and how animals switch from impulsive use to compulsive use despite the negative tradeoffs. This question requires a proper context where there is some tradeoff between drug use and life goals and survival. For example, phenomena in addiction include the reduced ability to maintain self-control and achieve goals; not everyone who first uses an opioid develops these impairments. The DSM-V defines aspects of this tradeoff in terms of human obligations (e.g., responsibilities to family and work)^28^; however, there is no comparable, ethologically relevant, and readily quantifiable task for rodents^29^. Neuroscience can now pursue naturalistic behaviors in the lab^30^, enabled by methods in computational ethology such as markerless pose estimation^31–33^. These methods enable assessing animal behavior in larger, less constrained environments with opportunities for richer decision-making.

We address these questions in two sets of behavioral experiments. First, we developed an assay where we genetically encoded iFentanylSnFR to record brain fentanyl levels alongside a machine vision routine to quantify behavior. To our knowledge, this recording is the longest continuous measurement of the brain [drug] alongside behavior (4 hours) and a first for drugs of abuse. We found a stereotypic, repetitive motor pattern that tracked the entire fentanyl time course. This pattern involved Straub tail, in which the tail curls up over the body, and hyperactive circling interrupted by stalling in the corners of the arena. This rise and fall of this effect correlate well with the rise and fall of the fentanyl concentration ([fentanyl]) over ∼3 h. This result challenges current models of cellular desensitization and acute tolerance that occur on ∼10 min and ∼1 h timescales, respectively. Second, to study the locomotor effect beyond the acute phase in an ethologically relevant survival task, we developed a paradigm based on foraging for water through a labyrinth maze^34^. Like in the open arena, mice in the maze receiving the same dose exhibited circling/stalling for ∼3 h, to the complete exclusion of successful foraging for water. Critically, this paradigm offers a normative definition of a deficit, as mice ought to have a baseline level of successful foraging to survive. We introduce this task to the substance abuse field as an additional metric for the deficits caused by opioid administration. Finally, we use these biosensors as purified protein and lyophilized powder to detect opioids in solution and human biofluids, enabling the engineering of continuous drug monitors in humans.

## Results

### A general approach to detecting opioids using OpuBC mutants

A common strategy in generating optical biosensors involves merging a conformationally switching protein with circularly permuted GFP (cpGFP) so that ligand binding elicits an increase in fluorescence^35–41^. The *µ*-opioid receptor could be a natural choice for detecting opioid drugs; however, unlike several other GPCRs, the *µ*-opioid receptor merged with cpGFP has been only minimally responsive to its ligands^42^. Periplasmic binding proteins (PBPs) have served as a conformational switch in genetically encoded biosensors such as iGluSnFR. PBPs are attractive for their stereotypically large Venus flytrap motion upon ligand binding. However, unlike neurotransmitters, neural drugs do not have cognate PBPs found in nature. We previously reported a biosensor for S-methadone based on this scaffold and a method of screening other drug classes towards new iDrugSnFRs^19,43–45^.

We sought to generalize from nicotinic biosensors based on a bacterial **<u>p</u>**eriplasmic choline-**<u>b</u>**inding **<u>p</u>**rotein (PBP), OpuBC (**Figure 1A and 1B, PDB: 7S7V**) to develop the iOpioidSnFRs in this work. We first tested the hypothesis that the binding pocket could be adapted to various opioids regardless of their activity at the *µ*-OR (e.g., agonists, partial agonists, and antagonists). We previously reported crystal structures of nicotinic ligands binding their optimized biosensors. Ligand binding to the PBP results in a conformational change, causing a glutamate residue in the 1^st^ linker to withdraw from quenching GFP’s chromophore. This allosteric linker-GFP mechanism appears agnostic to the binding pocket residues and ligands^19,44,45^. Docking simulations indicated that iNicSnFR3a’s binding pocket could accept opioids. The top-scoring poses placed the protonated tertiary amines of some opioids, such as tapentadol and levorphanol, where they could participate in cation-π interactions analogous to those between nicotinic ligands and their optimized biosensors^45,46^ (**Figures 1D and 1E**). Based on these results, we formulated a strategy to generalize OpuBC mutants toward various clinically used opioids (**Figure 1F**). We measured dose response relations between 28 opioids and nine biosensors, all based on OpuBC and previously reported for detecting nicotinic ligands (‘v4.6’-’v9’), ketamine (‘AK1’)^43^, and S-methadone (‘pre-Smeth1’) (**SI Figure 1, SI Table 1**).

ΔF/F_0_ = (F_sensor+drug_ – F_sensor_)/(F_sensor_) was calculated at each dose, and the low-concentration, linear portion of each dose response was fit with a linear regression. We have previously defined the δ-Slope for fluorescent biosensors as the slope of that linear regression with units *µ*M^-1^. Higher δ-Slopes are achieved by increasing ΔF_max_/F_0_ and lower K_d_. The δ-Slope expressed as a percent change *µ*M^-1^ for the library screen is visualized in shades of green, presenting at least an appreciable response to nearly all opioids (**Figure 1G**). A principal component analysis of the structures of the opioids’ Morgan fingerprints revealed clustering into the conventionally recognized opioid structure classes (**Figure 1H**). Each class displayed the greatest affinity for one or two biosensors. The monoamine opioids, such as tapentadol, showed structural similarity to nicotine and were best detected by iNicSnFR3b. v9 (iAChSnFR) displayed the strongest responses among members of the morphine and morphinan families. ‘AK1’ represents a distinct branch of the nicotinic biosensors, because the Y357G mutation at a critical cation-π residue ablates all responses to nicotinic ligands. ‘AK1’ uniquely detects fentanyl and its analogs with a modest S-Slope of ∼0.2 for fentanyl, which shares a comparable arylcyclohexamine motif to ketamine. Notably, a precursor to iS-methadoneSnFR removed by one point mutation was not the best hit for any of the 28 opioids. This result suggests that the diversity of its parent biosensors, not sensitivity to a particular opioid, provides a more effective search space for another opioid.

Using this mapping, we selected biosensor starting points and performed directed evolution in *E. coli*. The biosensors are soluble and retain their dose-response relations in bacterial lysates, enabling facile screening. Because the sensors typically required improved sensitivity, dynamic range, and selectivity, we mutated sites throughout the biosensor sequences. We performed iterative site-saturation mutagenesis, where we typically screened two to four positions per sensor and opioid pair and selected the mutant displaying the greatest increase in S-Slope to take forward. Remarkably, the OpuBC fitness landscape has been smooth enough for this approach and we have not yet encountered any “dead-ends” where the response could not be further improved. Examples of evolved leads for various clinically used opioids and one κ-selective opioid demonstrate the detection of their drug at ∼2 *µ*M (**Figure 1I**). Given that there is some mapping between OpuBC mutant sequences and opioid structures, we can narrow the search space to generate any next opioid sensor.

### Improving S-Slope by ∼500x through iFentanylSnFR1.0 and 2.0

Given the immediate human health relevance of detecting fentanyl, we then sought to take one of the weakest hits and generate the most sensitive iOpioidSnFR to date. The best biosensor hit for fentanyl, AK1, presented a relatively weak response with an S-Slope of 0.023 *µ*M^-1^. This hit is ∼3000x weaker than the best hit for tapentadol. Additionally, fentanyl’s six rotatable bonds and largely hydrophobic surface with few functional groups make it a challenging ligand for binder design^47^. Fentanyl also poses a unique challenge for the putative binding pocket interaction in OpuBC because its tertiary amine is placed near the middle of its longer, more linear structure compared to other drugs we have screened. The docked pose in iNicSnFR3a, where fentanyl’s protonated amine is deepest in the binding pocket, places the nitrogen ∼6 Å (upper limit for cation-π interactions) from aromatic residues and the attached carbons farther away and/or directed away from the aromatic residues. Furthermore, the amine’s substituents are completely removed from the perpendicular axis through the sidechain’s aromatic rings, likely not allowing cation-π interactions (**Figure 2A**). Only ‘AK1’, bearing a Y357G mutation which renders it insensitive to nicotinic ligands, displayed sensitivity towards fentanyl, indicating the need first to reshape the binding pocket.

**Figure 2:**
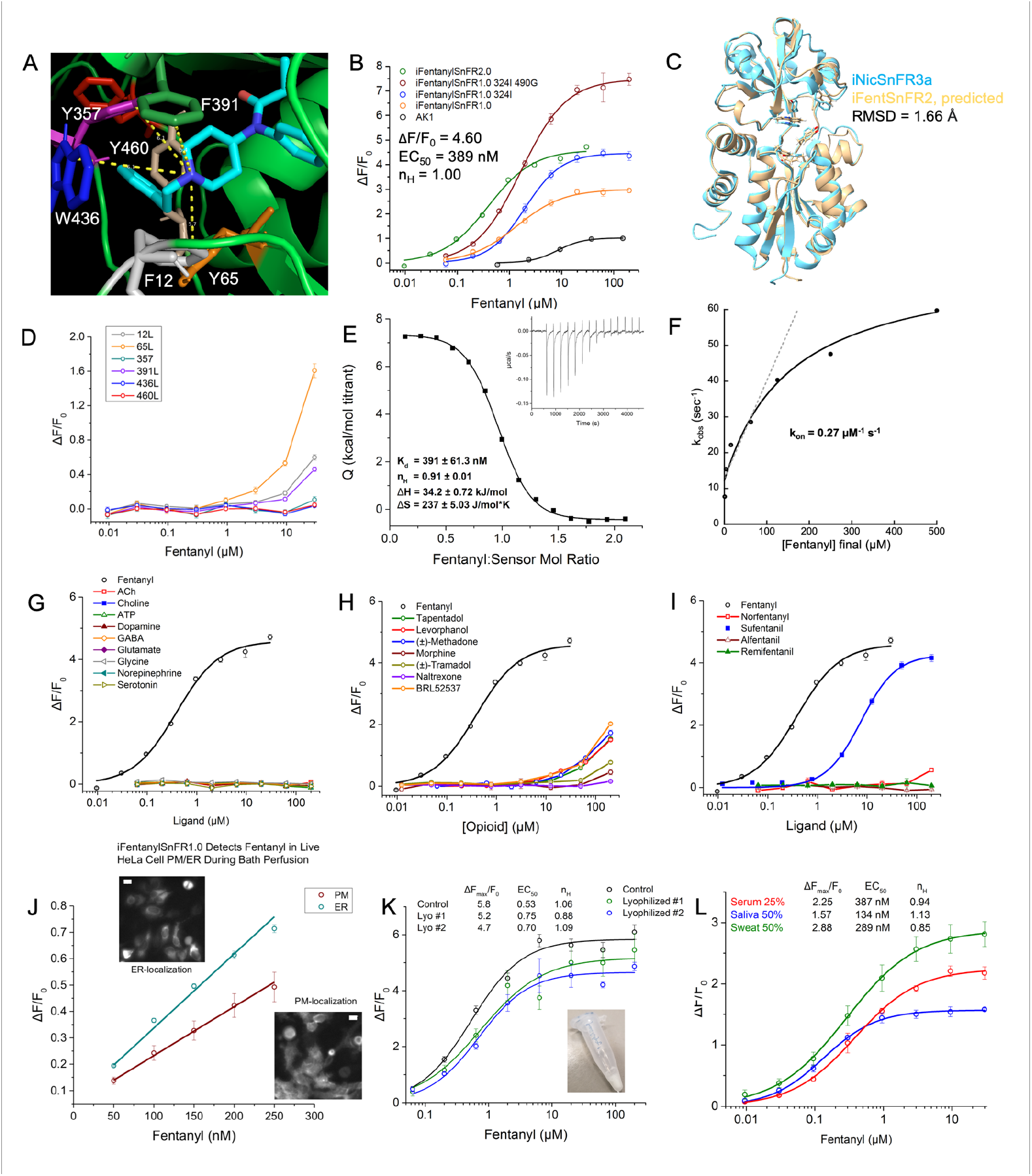
iFentanylSnFR evolution, characterization, and application in cells and biofluids. (A) Docking fentanyl in iNicSnFR3b (PDB: 7S7V). The top-scoring conformation shows fentanyl’s phenyl ring directed into the pocket, preventing full insertion of the tertiary amine into the aromatic box. (B) Directed evolution of iFentanylSnFR2.0 via, AK1, the initial hit bearing the founding Y357G mutation. (C) Overlay of the iNicSnFR3b PBP crystal structure (PDB: 7S7W, cyan) and the AlphaFold2 prediction of iFentanylSnFR2.0’s PBP structure (wheat). RMSD = 1.66 angstroms. (D) Leucine mutagenesis scan through the aromatic box diminishes dose responses. (E) Isothermal titration calorimetry: 2.5 μL of 200 μM fentanyl was injected into a cell with 20 μM iFentanylSnFR2.0 at 300 s intervals. Raw heat (figure inset) and thermodynamic parameters and stoichiometry are given. (F) k_obs_ vs. [fentanyl] in a 1 s stopped-flow kinetic response experiment. The k_on_ was determined by a linear fit of the first four points. (G) Dose response against neurotransmitters demonstrates complete selectivity. (H) Selectivity against other opioids shows complete selectivity beyond their pharmacologically relevant concentration ranges. (I) Dose responses against other fentanyl analogs: 20x selectivity against Sufentanil based on S-Slope and complete selectivity against the other analogs and the major metabolite, norfentanyl. (J) iFentanylSnFR1.0 monitors fentanyl permeation in living cells. HeLa cells were transfected with iFentanylSnFR1.0_PM and _ER. Dose responses ranging 50-250 nM show a linear response in both compartments with the S-Slope within a factor of 2x of each. Widefield imaging (figure insets for _PM and _ER constructs), 40x, 1.0 NA objective, 470 nm excitation, scale bars = 20 *µ*m. (K) Simulated field of iFentanylSnFR1.0 324I 490G: the purified protein was lyophilized (figure inset), stored in the dark at room temperature and humidity for 3 weeks, reconstituted in solution, and used in a dose response (n = 2). (L) iFentanylSnFR2.0’s dose response in biofluids spiked with fentanyl. Final v/v for serum = 25%, saliva = 50%, and sweat 50% mixed into a biosensor solution prepared in 3x PBS pH 7.4.For all dose responses: SEM is shown as error bars (n = 3 dose responses averaged).

We evolved AK1 further with the most extensive SSM experiments of all iOpioidSnFRs, spanning the 2^nd^ shell, hinge, and linkers (**SI Figure 3**). Two second-shell mutations, KI10G and RG395A, yielded a biosensor with ΔF/F_0_ ∼1 at ∼1 *µ*M that we termed iFentanylSnFR1.0 (orange trace, **Figure 2B**). Additional mutations across the PBP and the 2^nd^ linker yielded a sensor with ΔF/F_0_ ∼1 at 100 nM and S-Slope = 11.8 that we termed iFentanylSnFR2.0 (black trace, **Figure 2B**). This evolution campaign represents a 513x improvement in S-Slope from the initial hit to iFentanylSnFR2.0.

Still, only seven mutations, a ∼2% difference in the PBP’s sequence identity, separate this biosensor from iNicSnFR3a. Notably, all accepted mutations in the PBP selected alanine or glycine, and the sole linker mutation converted a proline to leucine. iFentanylSnFR2.0 preserves the overall fold in the PBP structure generated by AlphaFold2 with an RMSD = 1.66 Å with respect to the iNicSnFR3a’s PBP crystal structure (PDB: 7S7W) (**Figure 3C**). We scanned leucine through the binding pocket aromatic residues and found that preserving aromaticity at each residue was essential to binding fentanyl (**Figure 2D**). This screen also yielded the W436L mutant, the “null sensor” used in later experiments *in vivo*. The G357L mutation also yielded a null sensor, suggesting that increasing flexibility and/or reducing steric bulk at this position is critical to binding fentanyl. This result is in accordance with the library screen showing that only AK1 detected fentanyl family compounds.

**Figure 3:**
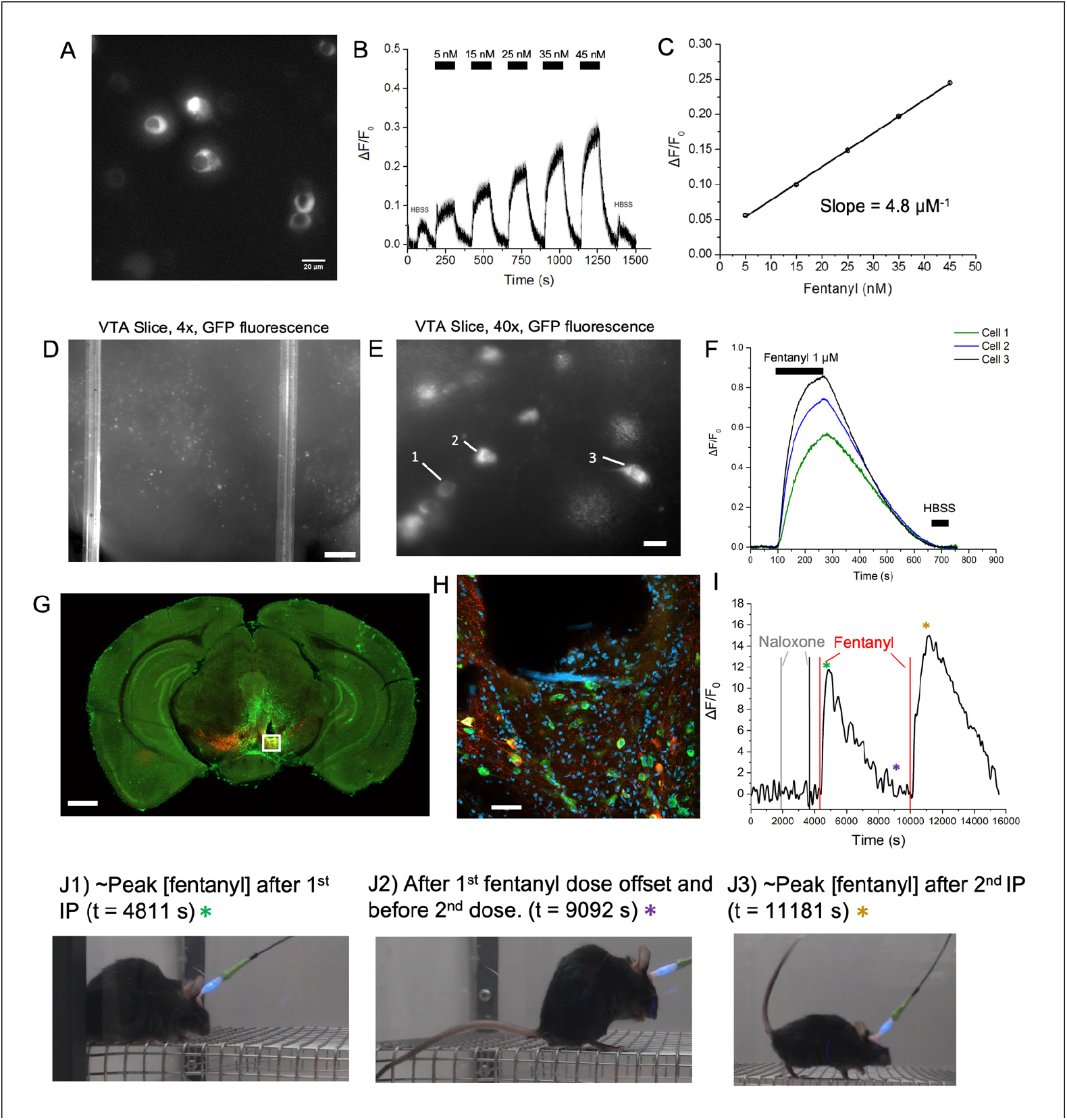
Detecting fentanyl in neurons enables real-time dosing in a freely behaving animal. AAV9-iFentanylSnFR2.0-cyto-WPRE was used for all experiments. (A-C): Linear dose response of iFentanylSnFR2.0 in primary hippocampal neurons. (A) Widefield fluorescence image (40x, NA = 1.0, 470 nm excitation). (B) The dose-response waveform during continuous imaging. A HBSS control (1 min wash in/wash out) was used to begin and end the experiment. Fentanyl applications (2 min wash in/wash out) were stepped by 10 nM from 5 nm to 45 nM. (C) Linear regression fit of responses corrected for the pH artifact (average of HBSS controls). (D-F): Imaging iFentanylSnFR2.0 in acute VTA slices. (D) Acute VTA slice (4x objective). Scale bar = 100 microns; brightness increased by 20%. (E) Individual cells show cytoplasmic targeting (40x objective). Cells in the focal plane used for analysis are annotated. Scale bar = 20 microns. (F) Waveform during the bath perfusion of 1 micromolar fentanyl (first arrow, t = 90 s) and washout (second arrow, t = 270 s). Bath exchange time ∼10 s. (G-H): Targeting the VTA in the intact mouse brain via fiber photometry – immunohistochemistry. (G) The coronal brain slice confirms fiber placement coordinates above the VTA. Tyrosine hydroxylase (Th, red) and GFP (green) channels are shown. Scale bar = 1 mm. The area below the fiber tract is denoted by the white box outline. (H) Zoom in on the fiber tract area (Th, GFP, and DAPI channels shown). Scale bar = 50 *µ*m. (I-J) Real-time dosing of fentanyl in an exemplar rodent via photometry. A mouse was pre-treated with 1 mg/kg and then 10 mg/kg naloxone (grey and black vertical lines). During the course of the first 1 mg/kg fentanyl dose (first vertical red line), no change in behavior was observed (J1, J2). The experimenter watched for the subsidence of the first dose and then administered the second fentanyl dose (second vertical red line). The animal then displayed the stereotypical Straub tail and circling behavior (J3). Asterisks match snapshots to time in the waveform in (I).

ITC determined a K_d_ = 391 ± 61.3 nM with a stoichiometry n = 0.91 ± 0.01 in agreement with the EC_50_ = 389 nM from the fluorescence dose response and the single binding site in the PBP (**Figure 2E**). iFentanylSnFR2.0 displayed an entropically driven binding interaction with its ligand like the parent cholinergic biosensors and other iOpioidSnFRs. Stopped-flow kinetics in a 1 s experiment determined a k_on_ = 0.27 *µ*M^-1^ s^-1^ (**Figure 2F**, raw data in **SI Figure 2C**). An extended measurement of iFentanylSnFR2.0 showed ∼95% and ∼100% of the maximum response ∼1 min ∼2.5 min, respectively, after the concentration jump from 0 to 1 *µ*M (**SI Figure 2F**). Accordingly, we later analyzed the photometry data with filtering with similar time constants.

iFentanylSnFR2.0 exhibits exquisite selectivity: no detectable response to any neurotransmitter or endogenous opioid peptide (**Figure 2G** and **SI Figure 2F**) and ∼zero response to other opioid drugs at ∼3 *µ*M and below, more than inclusive of their pharmacologically relevant ranges (**Figure 2H**). iFentanylSnFR2.0 discriminates even among its major metabolite, norfentanyl, and its analogs, sufentanil, alfentanil, and remifentanil (**Figure 2I**). Whereas the *µ*-OR displays an IC_50_ ∼3x lower for sufentanil than fentanyl ^48^, iFentanylSnFR2.0 displays a ∼20x greater response to fentanyl in terms of S-Slope.

As we evolved iFentanylSnFR, we found all candidates functional and applied some in various biological preparations. HeLa cells were transfected with iFentanylSnFR1.0_PM and _ER, directed to the plasma membrane (PM) and endoplasmic reticulum (ER), respectively (**Figure 2J**). Fentanyl equilibrated in ∼seconds in each compartment, and the S-Slopes were within a factor of 2x within each compartment, indicating that fentanyl rapidly permeates cells. In a simulated field test, samples of the predecessor to iFentanylSnFR2.0 were lyophilized and stored in the dark at room temperature and humidity for three weeks. The samples were reconstituted, and their dose response maintained at least 60% of the S-Slope and at least ∼80% of the dynamic range compared to a flash-frozen and thawed solution. Finally, in diluted pooled serum, saliva, and sweat spiked with fentanyl, iFentanylSnFR2.0 maintained its S-Slope within a factor of 2x. The responses differed primarily in their dynamic range beyond 1 *µ*M [fentanyl]. Filtrates, like sweat, provide a superior background for optical measurements.

### iFentanylSnFR2.0 in HeLa cells, primary neurons, acute slices, and the living murine brain

We transduced biological preparations with AAV9-hSyn-iFentanylSnFR2.0-cyto-WPRE to monitor fentanyl. Since fentanyl rapidly equilibrates across membranes, the cytoplasm affords an imaging volume suitable for PK. Additionally, the cytoplasm provides superior volume for imaging and does not require any specialized localization sequences. Primary hippocampal neurons were transduced with the viral construct and imaged under bath perfusion (2 min drug, 2 min wash) (**Figure 3A, B**). A sub-EC_50_ dose response ranging from 5 to 45 nM displayed a linear fit with S-Slope = 4.8 *µ*M^-1^ (**Figure 3C**). A full dose-response ranging from 30 nM to 3 *µ*M in ascending steps of √10 fitted to a Hill equation showed EC_50_ = 104 nM with ΔF_max_/F_0_ = 0.97 (**SI Figures 4A-C**). Both measures of the S-Slope demonstrate that iFentanylSnFR2.0 maintains its sensitivity within a factor of 2x in the primary culture preparation.

We chose to measure fentanyl waveforms in the VTA for the region’s central role in rodent models for substance abuse disorders, involved in the rewarding and locomotor effects^49,50^. In acute VTA slices, iFentanylSnFR2.0 exhibited widespread transduction (4x view, **Figure 3D**) and cytoplasmic localization (40x view, **Figure 3E**). The biosensor exhibited a similar response magnitude to a bath application of 1 *µ*M fentanyl as in primary culture. The imaging focal plane was 3 to 4 layers into the slice, ∼ 50 *µ*M from the surface. In this case, the biosensor’s response takes ∼3.5 min to plateau compared to ∼2 min. in the primary neuron culture (including glial cells).

We injected the viral vector in the VTA of live mice, followed by the implantation of a fiber optic element. After all experiments, immunohistochemistry confirmed fiber targeting the VTA and cytoplasmic localization iFentanylSnFR2.0-cyto. A representative coronal slice shows the fiber tract ending above the VTA with the characteristic bands of dopaminergic neurons visible in the midbrain (**Figure 3G**). A magnified region below the tract shows GFP-positive neurons partially colocalized with Th-positive neurons, as expected for pan-neuronal expression among midbrain dopaminergic and GABAergic neurons. The survival surgery was repeated with the null construct, AAV9-hSyn-iFentanylSnFR2.0-436L-cyto-WPRE. Photometry measurements in the animals after 1 mg/kg fentanyl IP showed no significant change (n = 3, **SI Figures 3D-F** and averaged gray trace in **Figure 4A** below). Therefore, any photometry signal is the result of fentanyl interacting with the binding pocket of iFentanylSnFR2.0 to elicit increased GFP fluorescence in the tissue.

**Figure 4:**
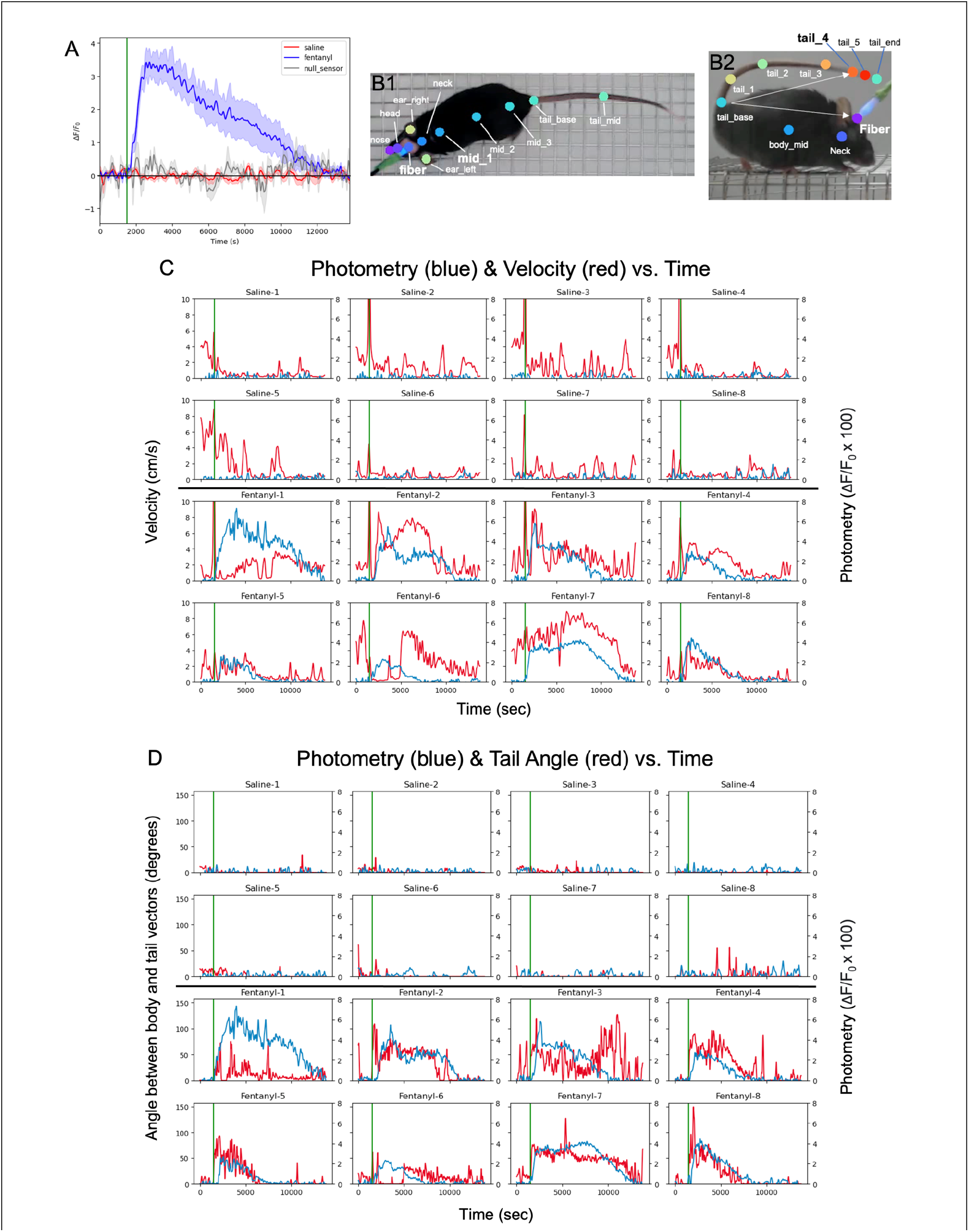
Recording [fentanyl] in the VTA Alongside Behavioral Quantified by a Machine Vision Routine. The photometry method prioritizes response kinetics over absolute quantitation and in situ recording, compatible with behavioral measurements. This focus serves our hypotheses regarding behavioral time course and acute tolerance. (A) Averaged photometry recordings of fentanyl in the VTA of freely behaving mice. Two cohorts expressing the functional biosensor received an IP injection of 1 mg/kg fentanyl (n = 8, blue) or saline (n = 8, red). A cohort expressing the null biosensor received an IP injection of 1 mg/kg fentanyl (dark gray). Waveforms are aligned to the time of IP (green line). SEM is shown as the shaded region. (B1-2) Machine vision tracking of mouse behavior designed by this work using top (B1) and side (B2) views. (C) Overlay of photometry waveform (blue, 60 s FFT filter) and the animal’s velocity (red) for each animal in the two cohorts: saline (top half) and fentanyl (bottom half). Velocity was calculated using a 3.33 s rolling window and was Gaussian smoothed (σ = 60 s). The vertical green line indicates the time of IP. (D) Overlay of photometry waveform (blue, 60 s FFT filter) and the animal’s tail angle elevation (red, Gaussian smoothed, σ = 60 s).

### Real-time dosing in an individual animal via photometry

We recorded from an animal while the experimenter observed and administered naloxone and fentanyl. iFentanylSnFR2.0 showed no response to naloxone and then, as expected, responded to the first fentanyl dose (1 mg/kg) with a relatively steep rise followed by metabolism/clearance (**Figure 3I**). During the majority of the first fentanyl waveform, the animal sat in a corner and did not display behavioral indications of fentanyl’s effects (**Figure 3J1**). Towards the end of the 1^st^ fentanyl time course, the animal showed a slightly elevated tail but still groomed itself, characteristic of a weak response to fentanyl (**Figure 3J2**). The experimenter watched for the complete subsidence of the fentanyl waveform in real time and then provided a second dose of 1 mg/kg fentanyl. At the 2^nd^ fentanyl waveform peak, the animal exhibited a stark Straub tail (elevated tail with a hunched, rigid body) and hyperactivity characteristic of *µ*-OR activation (**Figure 3J3**). Whereas naloxone is a pan-opioid competitive antagonist, fentanyl is a selective *µ*-OR agonist. This correlation between *µ*-OR agonism and the biosensor signal in the VTA encouraged a population study where the time course may vary between individuals.

### Population versus individual [fentanyl] waveforms

We then systematically quantified photometry alongside behavior in 16 animals in an open arena. The fentanyl treatment group consisted of eight animals and received a 1 mg/kg fentanyl IP injection; they displayed a significant positive waveform, whereas the eight animals that received saline showed no appreciable change from baseline (**Figure 4A**). The fentanyl treatment group average shows a ∼3 h time course. The population results agree with prior studies of fentanyl PK in mice. Conventionally, fentanyl pharmacokinetics has been performed by blood draws or tissue homogenization for a single time point analyzed by LC-MS. A previous study by Kalvass et al. sacrificed cohorts of animals at several time points and reported the population averages for blood and brain pharmacokinetics of fentanyl after a 0.9 mg/kg subcutaneous injection in mice. Kalvass et al. found that the [fentanyl] in the brain resembled that of serum within a ∼few min with a t_1/2_ of 4.9 min in the brain^51^. The results from Kalvass et al. displayed a similar overall time course and time constants for each phase, as observed in this work^51^.

In the present study, individual animals displayed time courses ranging from ∼2 h (e.g., animal Fentanyl-5) to ∼3.5 h (e.g., Fentanyl-1). This variability is evident in the population average, where the standard error is the largest mid-way through the subsidence of the waveform (**Figure 4A**). This result highlights the importance of measuring the relationship between drug PK and behavior within the same individual. Whereas microdialysis could be performed in situ, it is less robust at ∼nM concentrations and in detecting highly hydrophobic molecules^52^. Following the validation steps in prior biosensor studies, the stopped-flow and acute slice experiments in this work show that the biosensor resolves the PK in the animal.

### Machine vision approach to quantifying the behavioral effects

Cameras were centered above and on one side of an open rectangular arena, and we used DeepLabCut to train separate neural nets for each camera view using experimenter-labeled frames^31,32^. We labeled several more key points than required for the analyses as providing more labels in each frame improves training and increases the confidence of the predictions for any particular label^31^. In the top view, the animal’s midline, ears, and fiber were labeled (**Figure 4B1**). The “mid_1” point was used for the body’s displacement, and “fiber” was used for the head location. In the side view, the tail is extensively labeled in addition to the fiber and midline (**Figure 4B**). The “tail_base” and “tail_end” are bisected on the length of the tail by “tail_2”. “tail_3, 4, and 5” recursively bisect the tail down its length. C57BL/6 mice display variable colors at the end of their tails, and multiple key points were particularly useful in identifying the position of the tail. The tail angle is determined by the angle between the “tail_base” to the “fiber” vector and the “tail_base” to the “tail_4”.

The top view provides absolute quantitation of position and locomotor activity, with an average of 97% of all frames with sufficiently confident predictions of the “fiber” and “mid_1” positions (**SI Table 3**). 0.5% of all frames do not have the animals while they were handled for IP injection. The remaining ∼2.5% of other frames involve a subset of the animals in a collapsed position when they are inactive or during activities such as grooming and rearing such that the key points are not visible. In the latter case, the blocking of key points is brief (< 1 s). In sum, these ∼2.5% frames largely involve no movement. 93% of frames from the saline group video and 95% in the fentanyl group are accounted for where the tail data can be confidently rejected or accepted for analysis (**SI Table 4**). All videos were manually inspected to confirm that the time courses of the metrics matched the outputs from the machine vision treatment. Additionally, the cumulative key point location heat maps confirm a stark circling pattern in the top view (**SI Figure 5A**) and elevated tail position throughout the cage (**SI Figure 5B**) for the fentanyl group and not the saline group.

### The [fentanyl] waveform drives phasic locomotor activity and Straub tail

The photometry waveform for each animal is plotted against the velocity from the top view video (**Figure 4C**) and tail angle from the side view video (**Figure 4D**). The saline cohort provides a baseline for locomotor activity; these animals remain motionless in a corner, occasionally move to another corner, and occasionally groom. These animals display an elevated tail (acute tail angle) only incidentally, typically when their rears are pressed into a corner (**SI Figures 5A, B**). In contrast, the fentanyl treatment group shows phasic hyperactivity and Straub tail that tracks the fentanyl waveform. The beginning of this correlation is seen within a few min after IP injection (**SI Figure 6A**). Cross-correlations between each behavioral measure and the photometry waveform show a peak with a lag of 0-10 min for the fentanyl cohort except for the two animals that stalled and no appreciable correlation for the saline cohort (**SI Figures 6B and 6C**). Fent-1’s velocity drops to ∼zero at ∼1.5 h as it pauses in the corner. Fent-3 displays a high sensitization to the Straub tail effect; the first apparent phase is a typically elevated tail while circling, while the second phase tracks with the animal rearing frequently with a modestly elevated tail.

Fent-6 displays a ∼30 min period of inactivity and apparent low tail angle; however, inspection of the video shows that this individual collapsed from the fentanyl effect during this period. These exceptions warrant a finer characterization of the behavior.

### A stereotypical and paradoxical repetitive locomotor pattern

While hyperactivity, respiratory depression, and Straub tail have been observed after opioid agonism, short-term tolerance and how these effects interplay to produce overarching behavioral patterns are incompletely known. This work shows that “repetitiveness,” including those effects, appears to be the predominant, integrated effect. We define circling as the animal completing a circuit around the corners of the arena and the midpoints in between within 30 s. The saline cohort exhibits little or no circling: as animals largely move from one corner to another, where they spend most time sitting. In contrast, the fentanyl cohort displays phasic circling: the animal may spend up to ∼90% of a 5 min window circling, subsiding with the fentanyl waveform (**Figure 5B**). We define “nose in corner” as any period where the fiber optic entry point is within one cm of a corner, facing into that corner. This behavior is highly unusual insofar as prey animals prefer to stay close to the perimeter of arenas but facing outward. The saline cohort scarcely satisfies this criterion, whereas the fentanyl cohort dwells facing into the corner, sometimes standing on all four paws facing in and other times reared stationary, holding a Straub tail posture. After the fentanyl IP, the animals display up to ∼100% of their time faced into the corner in a 5 min bin (**Figure 5C**). In sum, the fentanyl cohort shows a significantly higher total distance traveled and time spent circling and with nose-in-corner than the saline cohort (**Figure 5D**).

**Figure 5:**
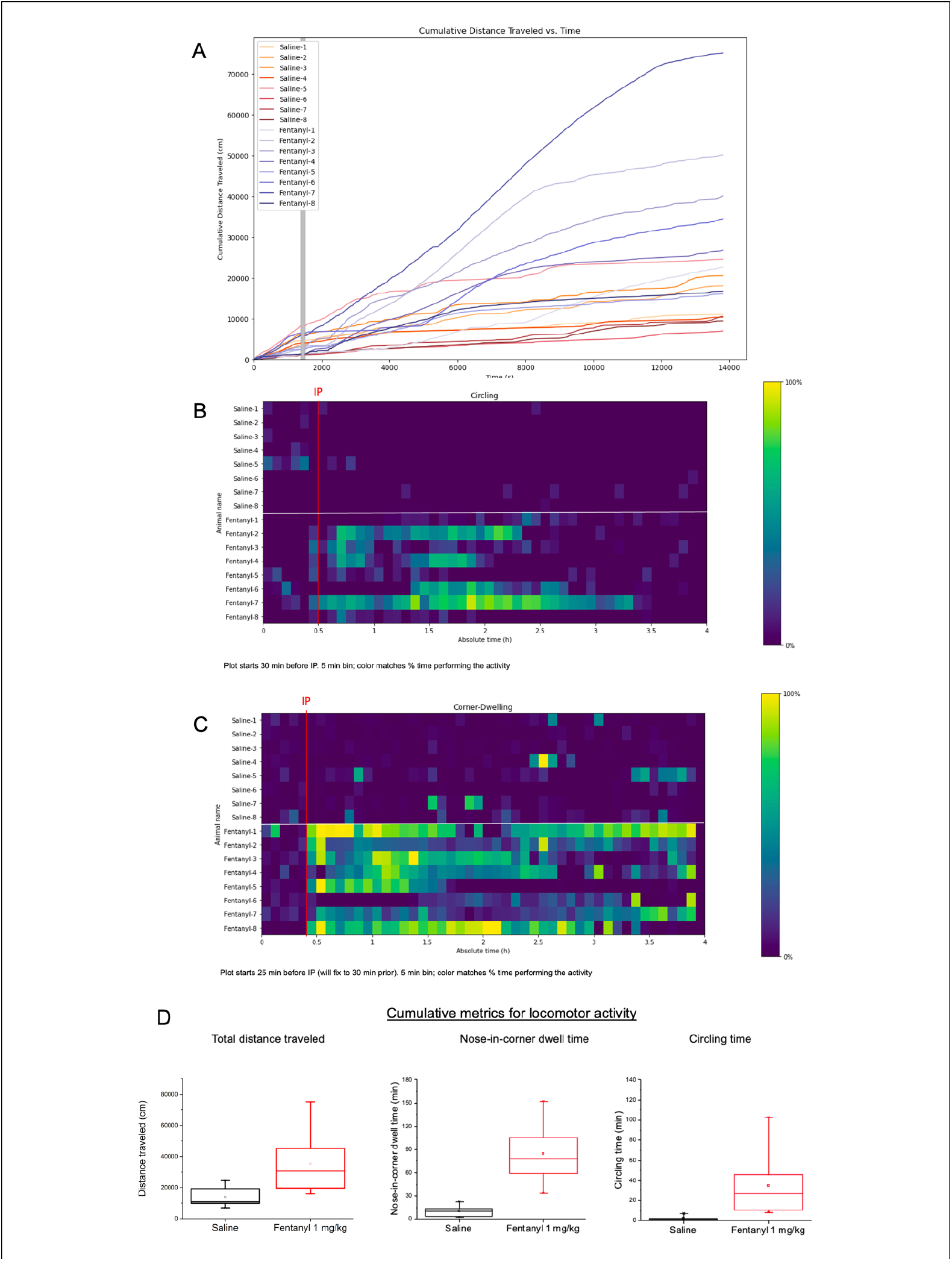
The stereotyped fentanyl effect demonstrated in cumulative statistics. (A) Cumulative distance traveled for each animal over the entire experiment (4 h). Individual traces are labeled for each animal. Time of IP indicated by vertical green line. (B) Time series heat map of circling behavior. Circling is defined as the animal passing through the center of the four sides of the arena within 30 s. Bins are 5 min. (C) Time series heat map of nose-in-corner behavior defined as periods where the animal places its nose into a corner of the arena continuously for at least 3 s. Bins are 5 min. (D) Comparison of cumulative locomotor behaviors between fentanyl (red) and saline (black) cohorts. The box denotes 25, 50, 75 percentiles and outlier whiskers denote the individual lowest/highest values.

We term the “fentanyl-driven stereotypical locomotor pattern” as a hyperactive repetitive pattern of circling, stalling, and Straub tail that follows the fentanyl time course. This pattern appears paradoxical in that animals on fentanyl show stalling by collapsing with visible respiratory depression or holding their “nose in the corner” but also display hyperactivity in terms of cumulative displacement. However, the apparent paradox is resolved when considering the time series data and the multiple circuits bearing *µ*-ORs. Although animals cannot simultaneously circle and dwell faced into the corner, they can display both behaviors within a 5 min bin. There is a phasic tradeoff between the rates of circling and dwelling with the nose in the corner. Based on the conventional view of neurocircuitry, a systemically administered *µ*-agonist would activate *µ*-ORs in the midbrain to elicit hyperactivity while also acting in the brainstem, inducing respiratory depression and a rigidified body as in the Straub tail phenomenon.

### Assessing the effect of fentanyl in an ethologically relevant survival task

We then tested the hypothesis that the stereotypical behavior due to fentanyl exposure is detrimental to self-maintenance assessed in a survival task. We chose a foraging paradigm where the animals learn to navigate through a labyrinth maze to find a water port (**Figure 6A**)^34^. Animals enter the maze on their own volution via a tube connected to their home cage. A single water port in the maze provides their only source of hydration (food is provided *ad libitum* in the home cage). Zero training is required for this paradigm, and the entire experiment is conducted during the animal’s dark cycle (active period) for 10 h. The maze consists of 128 numbered nodes, allowing for quantification and comparison of trajectories (**SI Figure 7A**). A camera placed below the maze’s IR-transparent bottom recorded the animal in the maze (**Figure 6B**). The maze setup and data analysis routine were reported previously in a study that showed animals exhibited “sudden insight” to learn direct routes to the water port^34^. In this work, one group of animals was deprived of water for 22 h prior to the experiment (“water-deprived”) and administered either saline (“Sal”), 0.1 mg/kg fentanyl (“Fent-low”), or 1.0 mg/kg fentanyl (“Fent-high”) IP at the start of the experiment (n = 8, each treatment). To assess intrinsic interest and navigation in the maze, another group of animals had food and water *ad libitum* in the home cage (“intrinsically motivated”, animal aliases given prefix “i-”) and was administered either saline (n = 6) or 1.0 mg/kg fentanyl (n = 4) to assess the impact of the drug on exploration in the absence of a foraging task.

**Figure 6:**
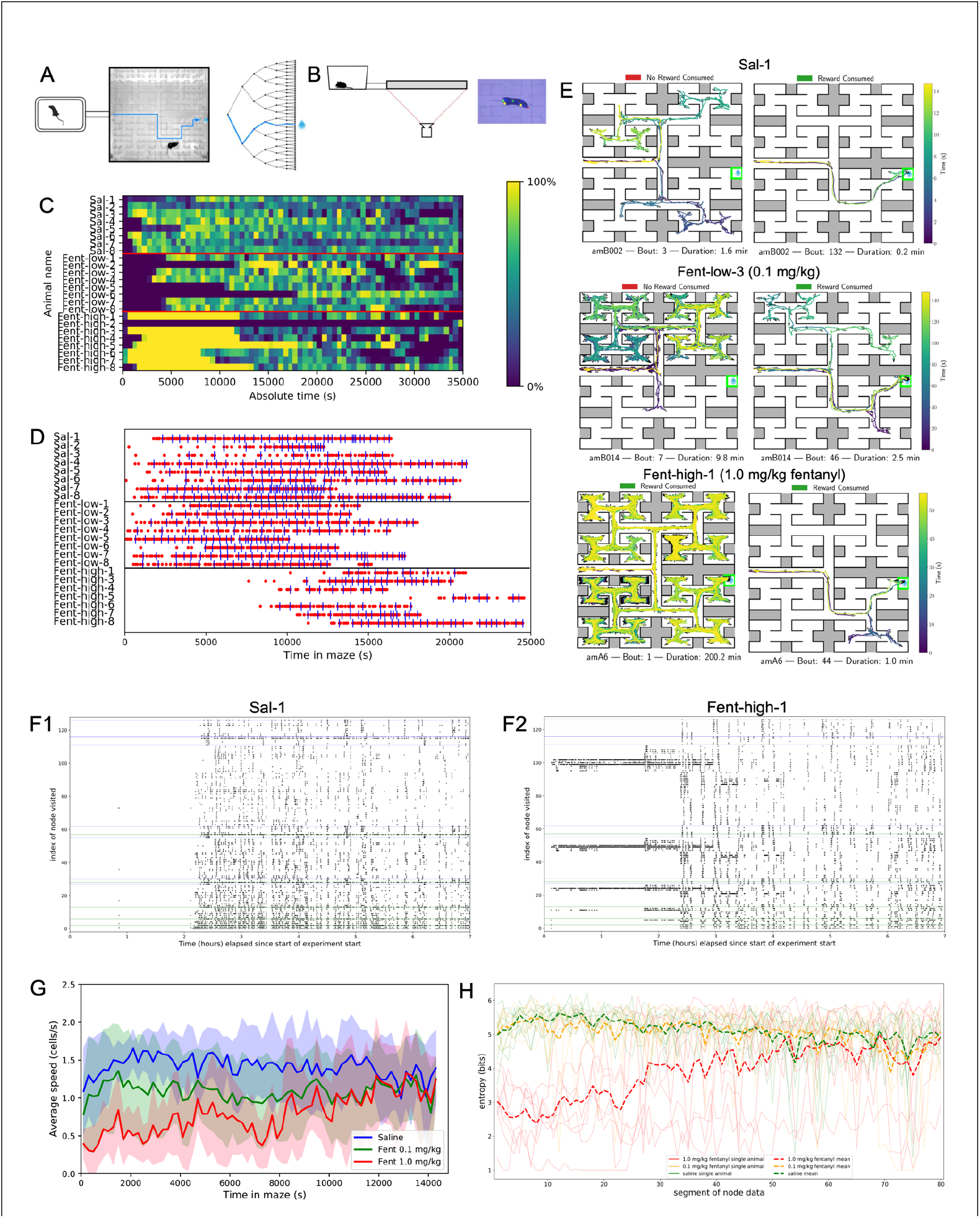
Fentanyl Effects on Animals Foraging for Water in a Labyrinth Maze. (A) Labyrinth maze paradigm with 2^6 binary decision branches and a single water port accessible at a terminal node. Mice were placed in a home cage with no water and ad. lib. food and could voluntarily enter the maze via a tube. The animals could earn 35 *µ*L water for nose-poking into a port at the end of the maze (90 s timeout after dispensing). The experiment lasted 10 h during the animal’s dark (active) cycle. (B) Videography scheme: the maze was constructed using IR-transparent plastic. An IR camera was placed underneath the maze to capture the animal’s movements, and the IR indicator light for water rewards at the water port. (C) Time series heat map for the occupancy of the maze over the entire experiment (0 to 100%, dark blue to yellow). Bins = 500 s. (D) Water rewards earned (red dot for each and blue tick mark for every 5^th^) over the cumulative time in the maze. amA7 was excluded from this analysis due to the first entry occurring at 9.5+ h. (E) Bouts from representative animals by fentanyl dose at early and mid-experiment time points. The path is traced as a line time series colored using a gradient from dark blue to yellow to represent the beginning and end of the bout. The duration of each bout is noted below the image. (F) Representative visualization of node visitation pattern for an animal that received saline (amB001, F1) or 1 mg/kg fentanyl (amA006, F2). (G) Velocity in the maze for saline (blue), 0.1 mg/kg fentanyl (green), and 1.0 mg/kg fentanyl (red) cohorts. The standard deviation is shown as a shaded region. (H) Entropy measure of each of the three groups of animals foraging for water in the maze (individuals are represented by solid traces and the group averages by dashed line).

The experimental groups displayed stark fentanyl dose- and time-dependent effects in navigating the maze and earning water. The water-deprived animals injected with saline showed immediate interest in exploring the maze and performed bouts typically < 5 min (**Figure 6C**). This group showed rapid (< 30 min) discovery of the water port and persistent successful foraging (**Figure 6D**). The water-deprived animals receiving 0.1 mg/kg fentanyl remained in their cage for 0.5-3 h before entering the maze. A side view of the home cage showed that these animals displayed the hallmarks of opioid agonism (e.g., circling and Straub tail) (**SI Video 3**). However, once these animals entered the maze at a later clock time compared to the saline group, they displayed rapid learning of the route to the water port. The water-deprived animals treated with 1.0 mg/kg fentanyl readily entered the maze and remained there for ∼3 h in a single bout. Animal ‘amA7’ is the exception as it entered the maze for the first time with ∼15 min remaining in the 10 h experiment. No water-deprived animal treated with 1.0 mg/kg earned water from the port for that first ∼2 h (wall or maze clock), and most animals only began earning water rewards 2.5 h or later into the experiment.

During their extended initial bouts in the water-deprived groups, the animals receiving 1.0 mg/kg fentanyl typically moved in a circling or figure-eight pattern in a 1/16 region of the maze, occasionally moving to another subsection but never finding the water port or drinking during this period. Notably, animals in this group will repeatedly press their face into a dead-end wall or corner, not immediately turning around. In contrast, animals receiving 0.1 mg/kg fentanyl display modest and brief or no such circling and stalling in the maze (after an extended period in the home cage – **Figure 6C**), and animals receiving saline never display impaired locomotion (representative traces, **Figure 6E**). The repetitive locomotion is characterized by revisiting a small number (< 16) of nodes without exploring other regions of the maze. This effect is visualized through representative raster plots of the nodes visited over time for an animal receiving saline versus 1.0 mg/kg fentanyl (**Figure 6F**). Another contrast in locomotor pattern between the groups is captured in part by the average speed over time. The 1.0 mg/kg fentanyl group initially displays ∼1/2 the speed of the saline and 0.1 mg/kg fentanyl groups while in the maze and eventually converge with these two groups after ∼3 h (**Figure 6G**). The repetitive circling locomotor pattern is also captured in a lower average entropy for the 1.0 mg/kg fentanyl group, converging with the saline group (**Figure 6H**).

In another set of experiments, the intrinsically motivated animals were treated with either saline or 1.0 mg/kg fentanyl IP and were allowed to enter the maze of their own volition. Animals receiving saline exhibited maze exploration behavior similar to the water-rewarded experiments (**SI Figure 7B-D**). As in the prior study establishing the maze paradigm, animals normally have an intrinsic interest in entering and exploring the maze thoroughly^34^. The animals receiving 1.0 mg/kg fentanyl also displayed similar locomotor and exploration results as in the water-deprived case: low entropy (**SI Figure 7B**) and repetitive circling (**SI Figure 7E**) despite extended residence in the maze (**SI Figure 7C**). 3 of 4 animals receiving 1.0 mg/kg fentanyl displayed extended bouts ranging from 2.5 to 4 h, and the 4^th^ animal did not enter the maze until 4 h into the experiment (**SI Figure 7C**), comparable to the outlier animal in the water-deprived, 1.0 mg/kg fentanyl treatment group. Therefore, the locomotor and exploration deficits from fentanyl administration appear to be independent of the water-deprivation and specific goal.

### Evolving selective sensors from pan-activating ligands: iTapentadolSnFR and iLevorphanolSnFR

Finally, we sought to evolve selective iOpioidSnFRs and fully characterize them for opioids with varying activities and pharmacokinetics. Given the biosensor library’s engineered responses to nicotinic ligands, particularly choline and acetylcholine, we sought to test the most challenging cases for improving selectivity. A principal component analysis of the S-Slopes from the “biosensor x opioid” library screen revealed that tapentadol, levorphanol, and meptazinol were outliers, given their strong pan-activation of the biosensors (**Figure 7A**). We then prioritized tapentadol and levorphanol because of their unique activities and pharmacokinetic profiles. Tapentadol, a *µ*-opioid agonist and norepinephrine reuptake inhibitor, is the most recent FDA approval for an opioid with a novel mechanism of action^53,54^. The combination of these two mechanisms allows for lower effective doses and has opened a route to less addictive opioids^55^. Levorphanol’s lack of cross-tolerance for *µ*-OR agonism with respect to prior morphine use^56^ and its longer duration of action (∼11 h half-life in humans) provide a pharmacokinetic basis for improved treatment of chronic pain^57^.

**Figure 7:**
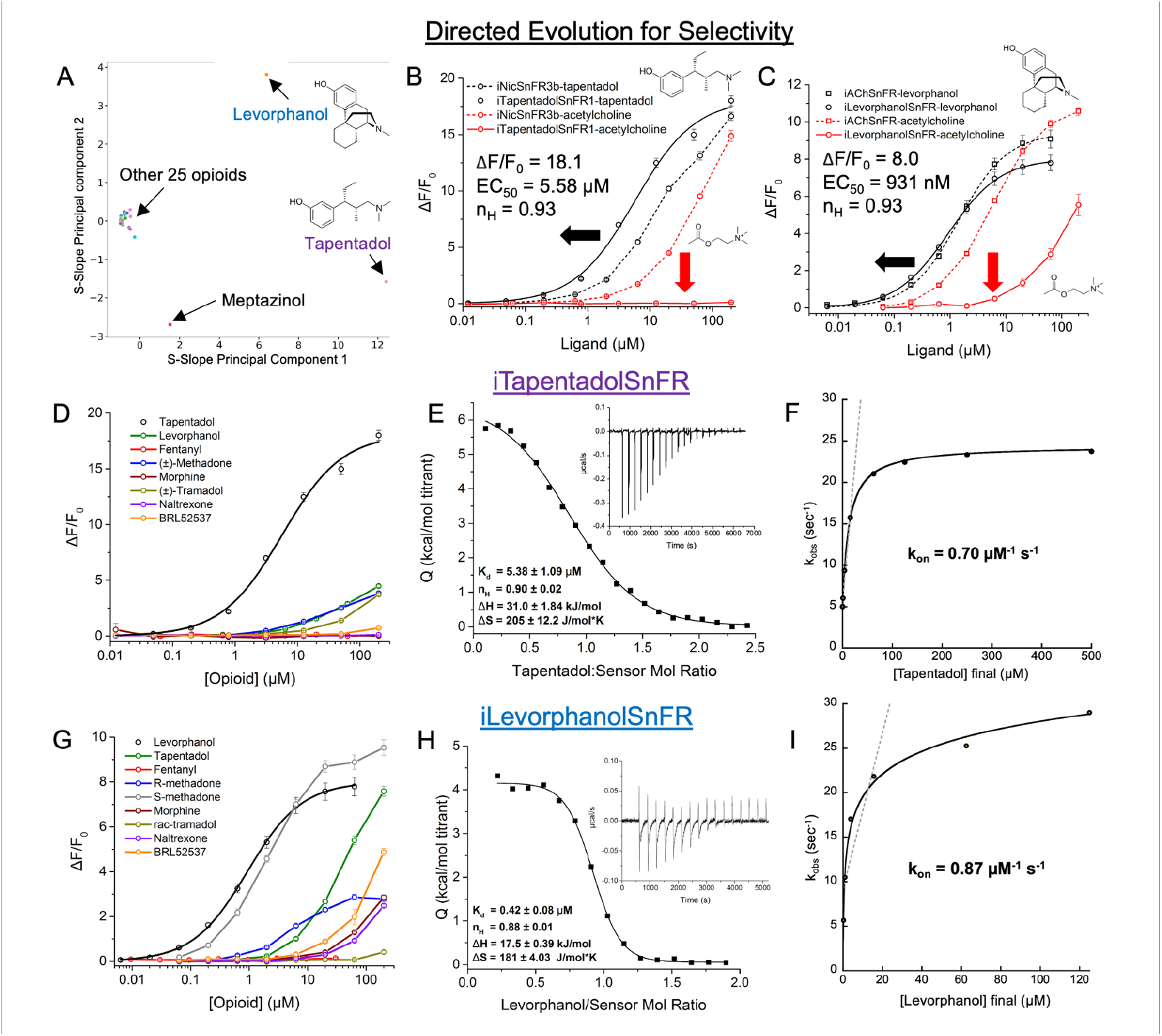
Evolving the two best hits for increased selectivity, yielding iTapentadolSnFR and iLevorphanolSnFR. (A) Principal component analysis of the cholinergic biosensor-opioid pair S-Slopes with outlier structures noted for tapentadol, levorphanol, and meptazinol. (B) A single point mutation, W436A, generates iTapentadolSnFR. Hill fit parameters for tapentadol against iTapentadolSnFR are listed. (C) Two point mutations in the 2^nd^ shell residues, QG15T and HA455P generate iLevorphanolSnFR and shift sensitivity to acetylcholine outside the physiological range. Hill fit parameters for levorphanol against iLevorphanolSnFR are listed. (D-F) iTapentadolSnFR characterization. (D) Selectivity against other opioid drugs show no response in their pharmacologically relevant ranges. (E) Isothermal titration calorimetry: 2 μL of 450 μM tapentadol was injected into a cell with 45 μM iTapentadolSnFR at 300 s intervals (raw heat, figure inset). Thermodynamic parameters given with fit error. (F) k_obs_ vs. [tapentadol] in a 1 s stopped-flow kinetic response experiment. (G-I) iLevorphanolSnFR characterization. (G) Selectivity against other opioid drugs demonstrated for all but S-methadone. (H) Isothermal titration calorimetry: 2 μL of 500 μM levorphanol was injected into a cell with 50 μM iLevorphanolSnFR at 300 s intervals (raw heat, figure inset). Thermodynamic parameters are given with fit error. (I) k_obs_ vs. [levorphanol] in a 1 s stopped-flow kinetic response experiment. For all dose responses: SEM is shown as error bars (n = 3 dose responses averaged).

OpuBC’s lack of affinity for endogenous ligands in mammals, save for choline, simplifies the search space for directed evolution. At most, only a positive selection for the opioid and a negative selection against acetylcholine/choline at each step were necessary. Additionally, we found that a small number of mutations was sufficient to engender the desired selectivity, further accelerating the approach to the desired biosensors. One mutation at a cation-π residue, W436A, in iNicSnFR3b, ablated all sensitivity to cholinergic ligands while improving an already strong response to tapentadol to yield iTapentadolSnFR (**Figure 7B**). iLevorphanolSnFR was generated via two mutations in the second shell, G15T and A455P, diminishing the sensitivity to ACh outside the physiologically relevant range (**Figure 7C**). Both variants show little or no response to other neurotransmitters (**SI Figures 2A, 2B**), and the endogenous opioid peptides, met-enkephalin, endomorphin-1, and *α*-endorphin (**SI Figures 2D, 2E**). iTapentadolSnFR displays ∼zero sensitivity to other opioids at ∼1 *µ*M and below, spanning their clinically relevant ranges. iLevorphanolSnFR displayed selectivity against all tested drugs except S-methadone (**Figure 7G**). iTapentadolSnFR and iLevorphanolSnFR are well-behaved insofar as their K_d_ values, determined by isothermal titration calorimetry (ITC), are within a factor of 2x of their EC_50_ from fluorescence with a stoichiometry near 1.0 (**Figures 7E and 7H**). Stopped-flow kinetics showed that iLevorphanolSnFR’s k_on_ is approximately unchanged from the k_on_ for its parent, iAChSnFR^58^, used to detect synaptic acetylcholine release *in vivo*. iTapentadolSnFR has an ∼order of magnitude larger k_on_ than its predecessor, iNicSnFR3a^43^ (**Figures 7F and 7I**, raw data in **SI Figures 2A and 2B)**.

In sum, we can generate sensitive and selective responses to opioids even if they display promiscuous binding to the starting points. Most notably, nine variants based on OpuBC provide enough diversity in starting points so that a small number of further mutations can yield the desired selectivity. These results point to “activity cliffs” in sequence space where small steps in the sequence landscape result in dramatic changes in function. Activity cliffs represent a weakness in current machine learning approaches to protein engineering; however, this weakness can be resolved by collecting training data like these, and including a training criterion assessing performance with activity cliffs^59^. These results suggest that the binding pocket and 2^nd^ shell residues in OpuBC harbor the potential for activity cliffs, exploited for converting selectivity from one drug class to another.

## Discussion

We report the first class of sensitive and selective genetically encodable biosensors of synthetic opioid drugs. These sensors enable PK studies spanning the subcellular to the whole animal. We focused on the relation between whole-body PK (as measured in the brain, specifically VTA) and rodent behavior to assess the effects on locomotor activity. That the stereotypical pattern tracks the entire fentanyl waveform was unanticipated: conventional models show desensitization and “short-term tolerance” on time scales of ∼10 min and ∼1 h, respectively^60^. In the ligand bias model of GPCR activity, fentanyl displays strong β-arrestin bias, and we expected it to induce strong, fast desensitization^60^. Instead, we observed that the fentanyl time course drove locomotor activity with similar kinetics ranging from 1.5 to 3 hours, well beyond the conventional regime of cellular desensitization. In a simple model, behavioral effects are directly mediated by neuronal dynamics which includes receptor kinetics and cellular allostatic mechanisms. This work calls for the reconciliation of the dynamics in cellular and circuit models with the observed behavioral and fentanyl time courses.

Additionally, our behavioral analysis points to the need for additional mechanistic study of the unique effects of opioids in rodents not seen in humans and dissection of the contributing mechanisms. While “high doses” of fentanyl would be normally associated with incapacity, in rodents, they elicit erratic hyperactivity. There is some similarity between the rigid posture of Straub tail that includes the mouse’s whole body and chest wall rigidity (first termed “wooden chest syndrome”) in humans^61^. However, to our knowledge, there are no reports of hyperactivity and repetitive locomotor behavior in humans. There is a monotonic relationship between increased fentanyl dose and increased respiratory depression^62^ and peak hyperactivity at 1.0 mg/kg fentanyl^63^, the dose used in this work. However, prior works had not reconciled the apparent paradox of Straub tail/respiratory depression alongside hyperactivity. This work shows that an individual animal administered fentanyl can display periods of hypo- and hyper-activity, switching every few ∼min, all while displaying the Straub tail. The alternation between hypo- and hyper-activity could arise from hypoxia-dependent compensatory activity, as in Cheyne-Stokes or Biot’s breathing (switching between periodic high- and low-breathing rates) during chronic opioid exposure^64^.

This work introduces a method to address a DSM-V criterion for substance abuse: dysregulation of behavior that impacts self-maintenance. Conventionally, the substance abuse field relies on tasks like the 5-choice serial reaction time task to assess the drug’s effect on an animal’s self-control^65^; however, this task requires weeks of training and a non-natural. In contrast, mice readily forage in complex environments for resources, both in the wild in and our experiments. That mice do not require any prior training whatsoever suggests that this navigation task assesses a core capacity of the animal that is necessary for survival. In this sense, the maze navigation task more closely matches the DSM-V criteria than conventional cognitive tasks. Indeed, we see dose-dependent aberrations from normal exploration and foraging, where mice in the 0.1 mg/kg fentanyl cohort circle in the home cage and enter the maze later compared to the saline cohort. The 1.0 mg/kg fentanyl cohort showed circling in a small portion of the maze for several hours. Additionally, the experiment lasts 10 hours, allowing for the subsidence of fentanyl, into a potential withdrawal period. Typically, experimenters precipitate withdrawal with an opioid antagonist or wait and observe later in an open arena. In contrast, the maze paradigm monitors continuously in a task with clear survival value, providing an integrated view of cognition. The animals show a rapid recovery in foraging performance after ∼3 hours and sustain it for the rest of the night; they are not completely debilitated during the first withdrawal period. Future studies will investigate repeated opioid exposure to determine if deficits are restricted to the period of the drug time course and the threshold at which repeated exposures lead to failure in survival tasks^27^.

Long-term tolerance is a critical feature of OUD, driving escalated intake and, potentially, death by overdose. This work’s continuous monitoring method suggests one future tactic to assess long-term tolerance: identify administration regimens that lead to behavior deviating from the fentanyl waveform. We know that other neural drugs, generally, including some other opioids, lack a straightforward relationship between free concentration and activity. For example, the morphine family of drugs has several active metabolites, buprenorphine has an unusually slow dissociation constant that is rate-limiting for the analgesic effect^66^. Other opioids, such as tapentadol, act on multiple receptors. In contrast, fentanyl offers straightforward pharmacology as it is not a prodrug, lacks active metabolites, and selectively binds to MOR versus all other receptors. With the present work, we have a straightforward mapping between [fentanyl] waveform and locomotor behavior.

Continuous monitoring devices to perform analogous studies in humans would allow researchers and drug abuse therapy providers to personalize dosing methods to minimize side effects, particularly in reducing tolerance. With advances in diagnostic device fabrication, the bottleneck in continuous monitors has moved to generating selective, sensitive, and robust sensor molecules paired with appropriate signal transduction mechanisms^67,68^. This work demonstrates a protein with excellent conformational switching ability with little or no affinity for the drug of interest can be evolved to become an optimal binder. At present, we believe that the OpuBC-based tactic will succeed for many drugs of MW < ∼700 capable of making cation-π interaction. These results suggest iFentanylSnFR2.0’s optical response can be calibrated to the biofluid, meeting existing form factors for both rapid point-of-care tests and continuous monitors comparable to a fluorescence-based continuous glucose monitor^69^. This result is particularly useful for addressing the proliferation of synthetic fentanyl analogs^70,71^ and research compounds such as unconventional *µ*-opioids and synthetic kappa opioids under consideration in therapeutic applications^72,73^. Using iLevorphanolSnFR alongside methadone maintenance therapy would require controlling for S-methadone using another biosensor. Otherwise, both iOpioidSnFRs are amenable to experiments or diagnostic readouts involving polypharmacy.

These tactics could be applied to other scaffolds that exploit the many naturally occurring and synthetic^47^ conformation-switching proteins. While genetically encoded sensors have been reported for many neurotransmitters, there remain many small molecule drugs and metabolites to monitor in situ for metabolic, cell signaling, and pharmacokinetic interests. Nature has afforded ∼25,000 bacterial ligand-binding proteins alone, and a recent work demonstrated a technique to screen for sites where one can insert reporters, including the present cpGFP^76^. PBPs offer excellent physical properties, including large (10-20 Å) conformational changes, aqueous solubility, stability as a lyophilized powder, and lack of crosstalk with mammalian cells. In contrast, GPCR-based biosensors are not soluble and stable as a powder which limits their application in subcellular PK (they cannot be directed to organelles) and most point-of-care diagnostics. Recent advances in signal transduction techniques, such as nanopores, will enable further applications of binding-activated conformationally switching proteins, including PBPs^76^. The protein engineering and computational ethology approaches together serve the goal of detecting molecules in living organisms as they behave in their ecological niches.

