## Supplemental Information for "Correspondence of fentanyl brain pharmacokinetics and behavior measured via engineering opioids biosensors and computational ethology"

### Supplemental Figures 1-7

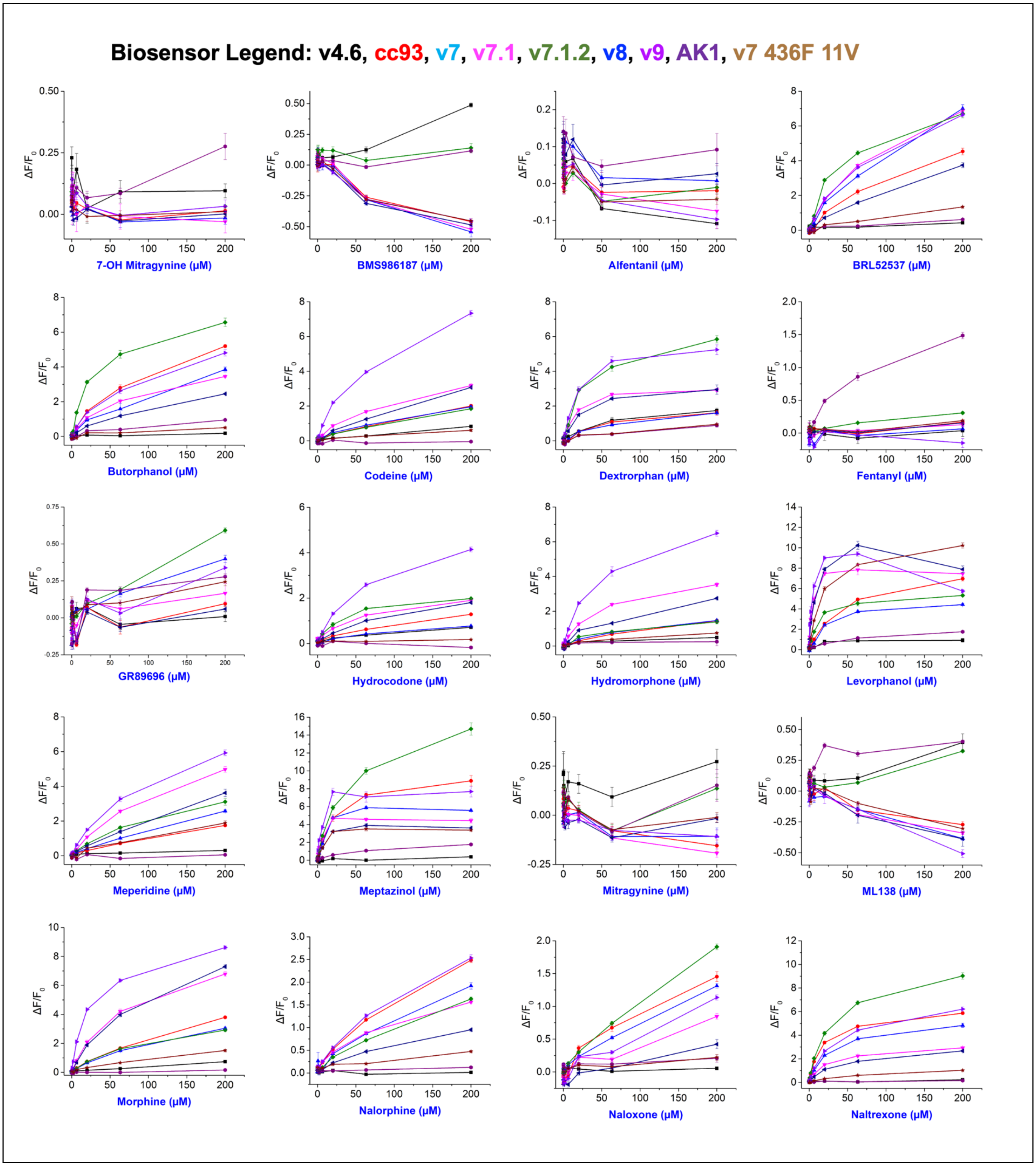

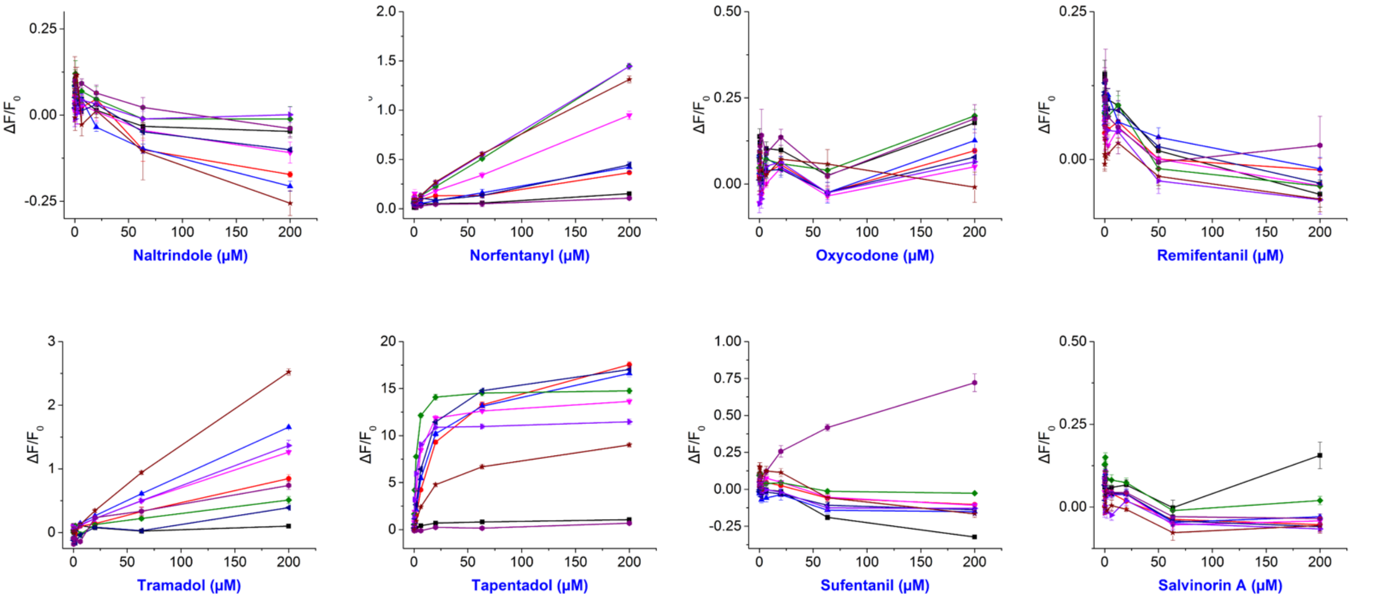

**SI Figure 1 related to Figure 1**: Raw data for biosensor responses plotted against each opioid ligand. A serial dilution of each opioid was prepared and mixed into a biosensor solution to achieve a final [opioid] ranging from 200 µM to 63.3 nM with a constant [biosensor] of 100 nM. Each dose response is shown as a line graph (color legend identifies biosensor sequence). These dose responses determined the linear range for the regression for each opioid vs. biosensor. The SEM is shown as error bars (n = 3 dose responses averaged).

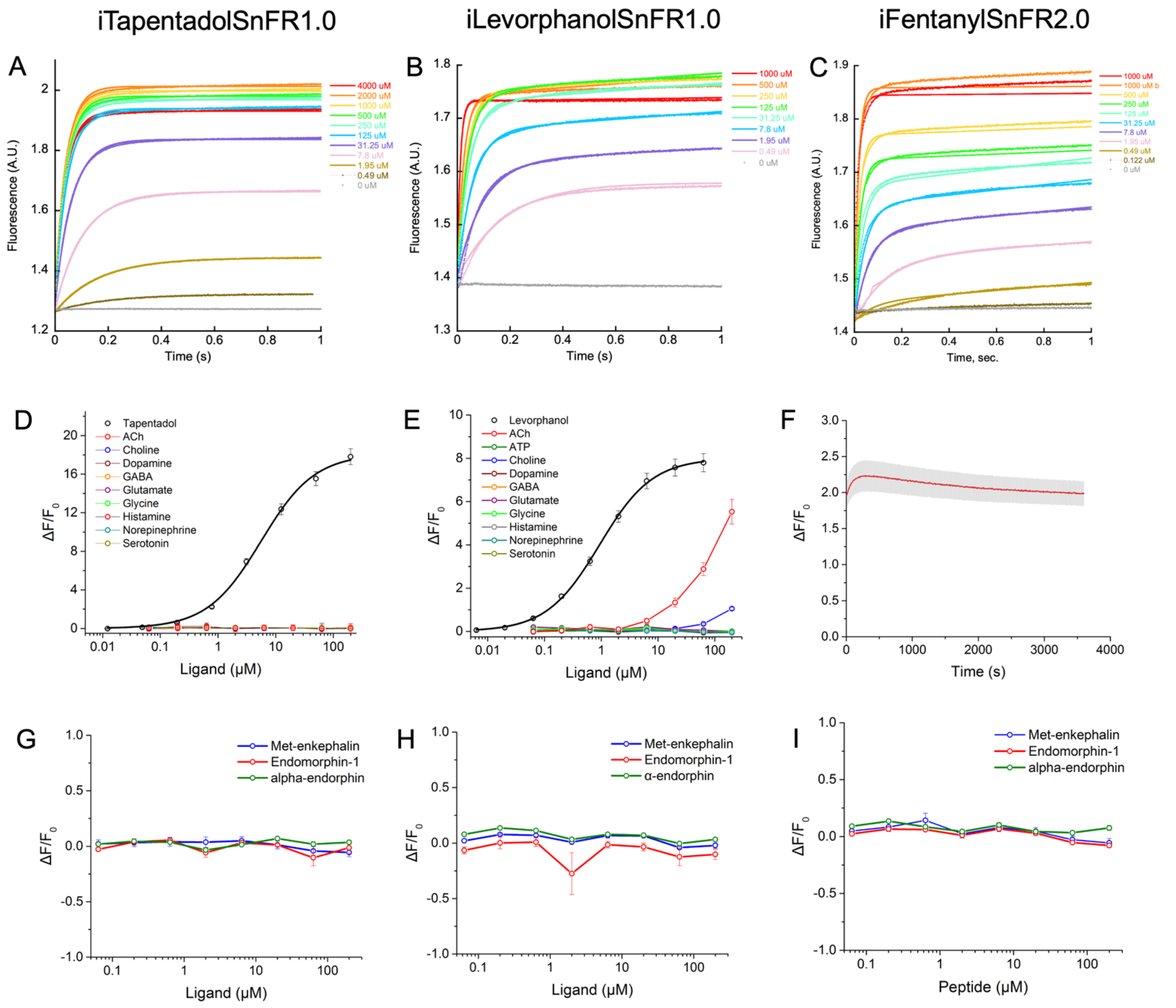

**SI Figure 2 for Figures 2 & 7**: Raw data and additional characterization of iTapentadolSnFR, iLevorphanolSnFR, and iFentanylSnFR2.0.
(A-C): 1 s stopped-flow kinetic data. 200 nM of each biosensor was mixed in equal volume to each ligand in varying concentrations as labeled. The final [opioid] in the chamber was one-half of this value.
(F) 1-hour measurement of iFentanylSnFR2.0's response to 1 $\mu$M fentanyl shows ~90% of the response immediately (~3 s) after mixing followed by a ~4 min equilibration for the remaining ~10% of the signal and then linear bleaching.
(D-E): Selectivity against neurotransmitters (iFentanylSnFR2.0's response shown in the main text).
(D) iTapentadolSnFR shows no response to any neurotransmitter, including acetylcholine, owing to a mutation in a critical cation-π residue, like iFentanylSnFR2.0.
(E) iLevorphanolSnFR shows diminished response to acetylcholine with S-Slope < 0.05 and ~zero response at the physiologically relevant concentrations (at ~2 μM and below).
(G-I): Selectivity against endogenous opioid peptides in fluorescent dose responses. No significant response is observed for [peptide] at the highest tested dose (200 μM) or below for any sensor.
For all dose responses: SEM shown as error bars (n = 3 dose responses averaged).

### SI Figure 3: iFentanylSnFR2.0 Directed Evolution Tree

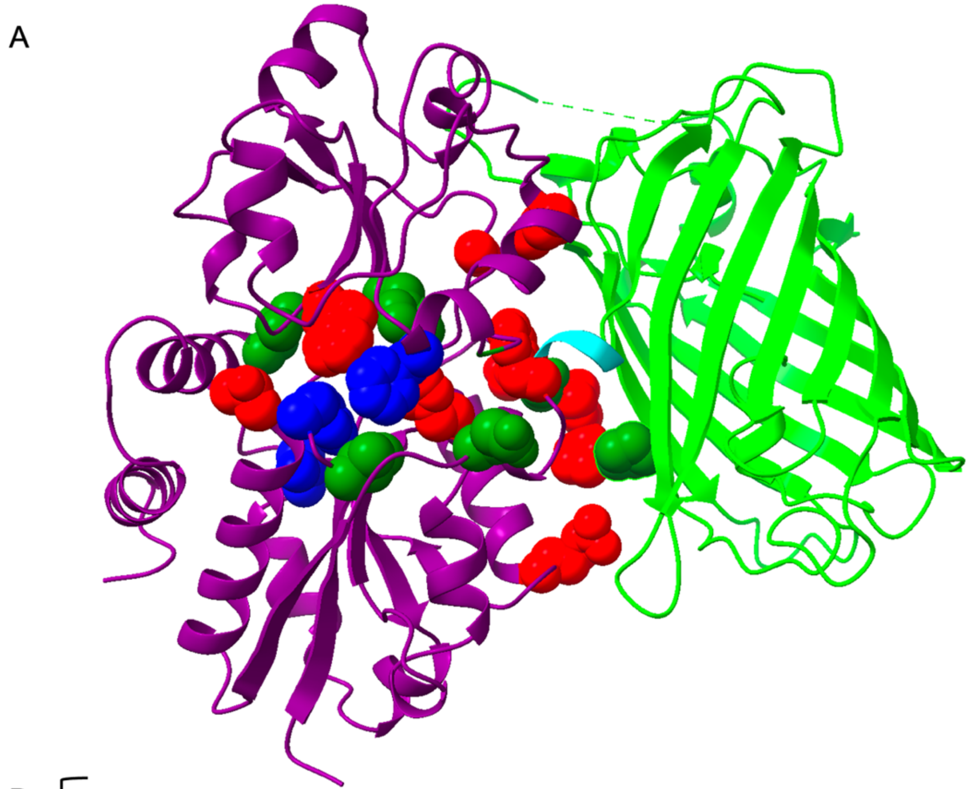

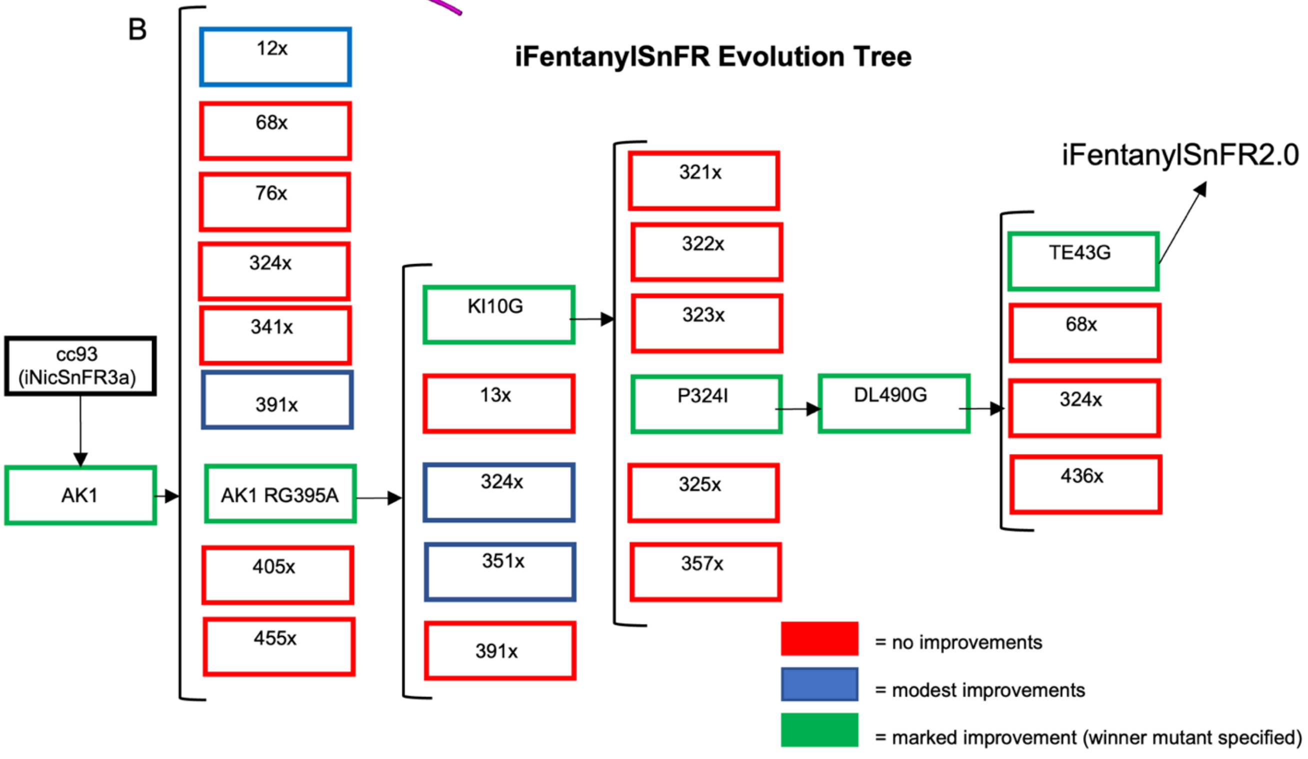

**SI Figure 3 for Figure 3**: iFentanylSnFR2.0 Directed Evolution Tree.

(A) Crystal structure of iNicSnFR3b (PDB: 7S7V) annotated with residues mutated in the directed evolution towards iFentanylSnFR2.0. Side chains mutated are shown as spheres: failed mutations (red spheres), modest improvements not taken forward (blue spheres), and accepted mutations (dark green spheres).

(B) Evolution tree representing site saturation mutagenesis experiments. Residue nomenclature: first and second residues before the position number are the amino acids in the OpuBC homologue from *Thermoanaerobacter sp* X513 in nature and iNicSnFR3b, respectively. The residue listed after the position number is the amino acid found in iFentanylSnFR2.0.

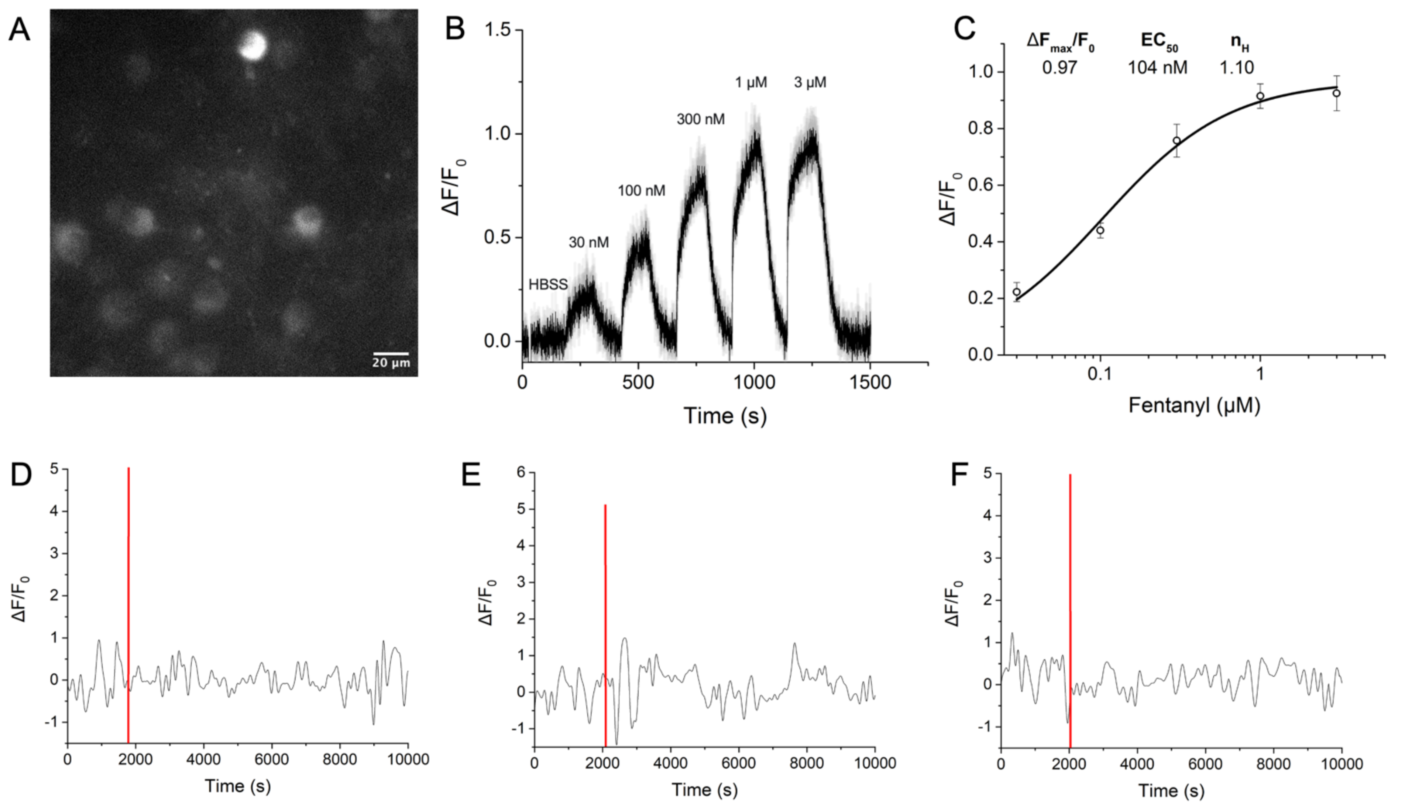

**SI Figure 4 Related to Figure 4 & 5.**

(A-C) iFentanylSnFR2.0 response in primary hippocampal neurons under saturating [fentanyl].

(A) Widefield image of the neurons transduced with iFentanylSnFR2.0 (scale bar = 20 microns). 40x objective, 1.0 NA, 470 nm excitation. (B) The waveform of biosensor response during widefield fluorescence imaging. The HBSS control and then increasing [fentanyl] bath application (2 min on, 2 min washout) was applied. (C) The steady-state response after correcting for the HBSS artifact was fit with the Hill equation (parameters shown). The dynamic range observed in primary cell culture is comparable to that of the acute slice response to 1 micromolar fentanyl bath perfusion.

(D-F) Individual traces from the "null" negative control experiment. iFentanylSnFR2.0 W436L (null sensor) was cloned into the same pAAV vector, packaged in AAV9, and injected into same coordinates in the VTA, replicating the protocol used for the functional sensor. Photometry recordings were conducted before and after administration (1 mg/kg fentanyl IP time noted by vertical red line). No appreciable response was observed in response to the IP.

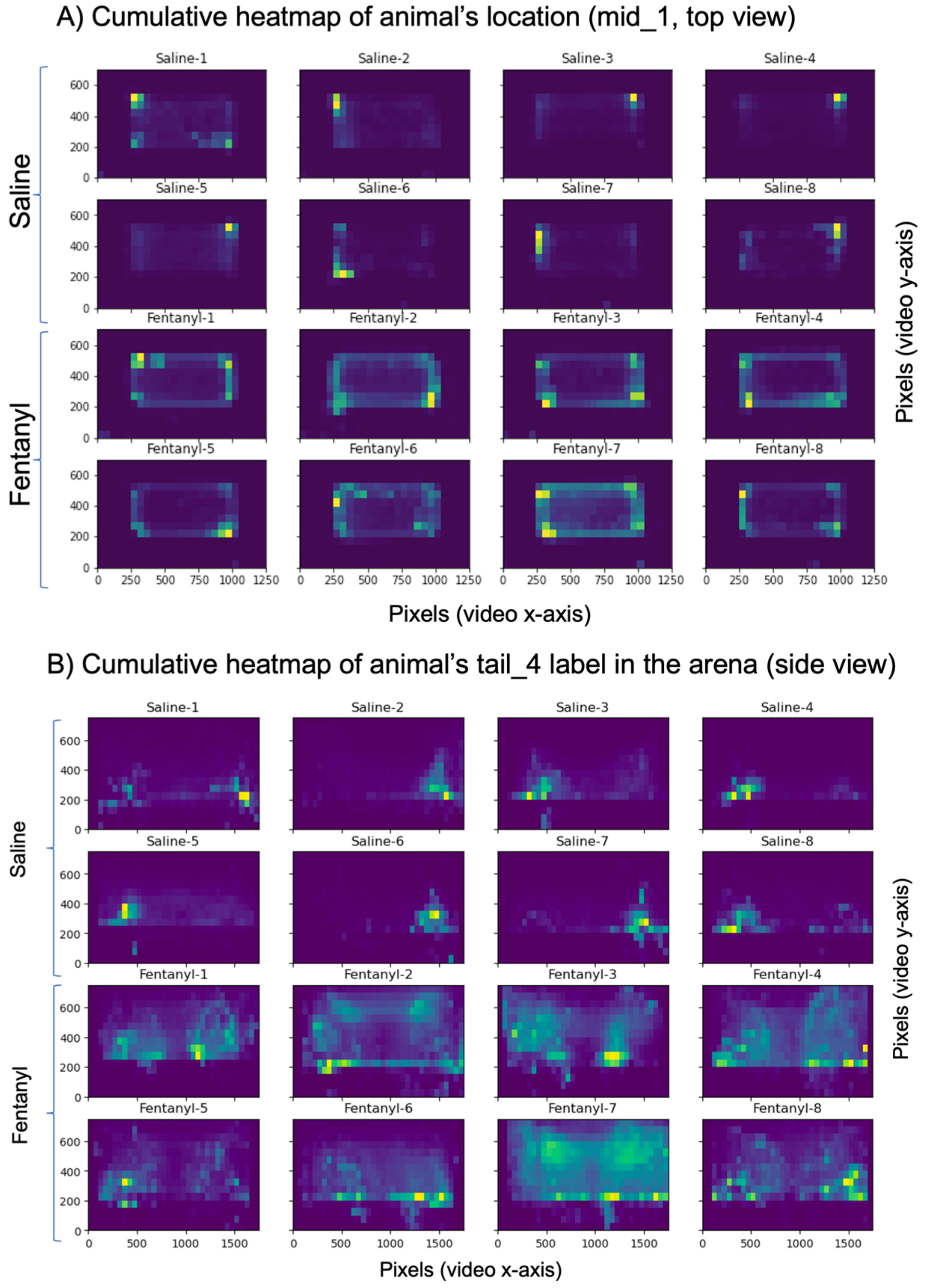

**SI Figure 5 Related to Figure 5 & 6: intermediate checks on raw machine vision data.**
(A) X-Y position heat map (top view) of the animal's posterior body label. Animals receiving fentanyl show a circling pattern whereas the saline cohort spends most time sitting/grooming in the corners of the arena.
(B) X-Z position heat map (side view) of the second to last label on the animal's tail ("tail_4", arrow in inset image of the cage). Animals receiving fentanyl show an elevated tail throughout the cage (i.e., moving around the arena displaying Straub tail). Animals receiving saline show the tail largely lies on the floor, hangs below the mesh, or is pushed up the wall when sitting in a corner.

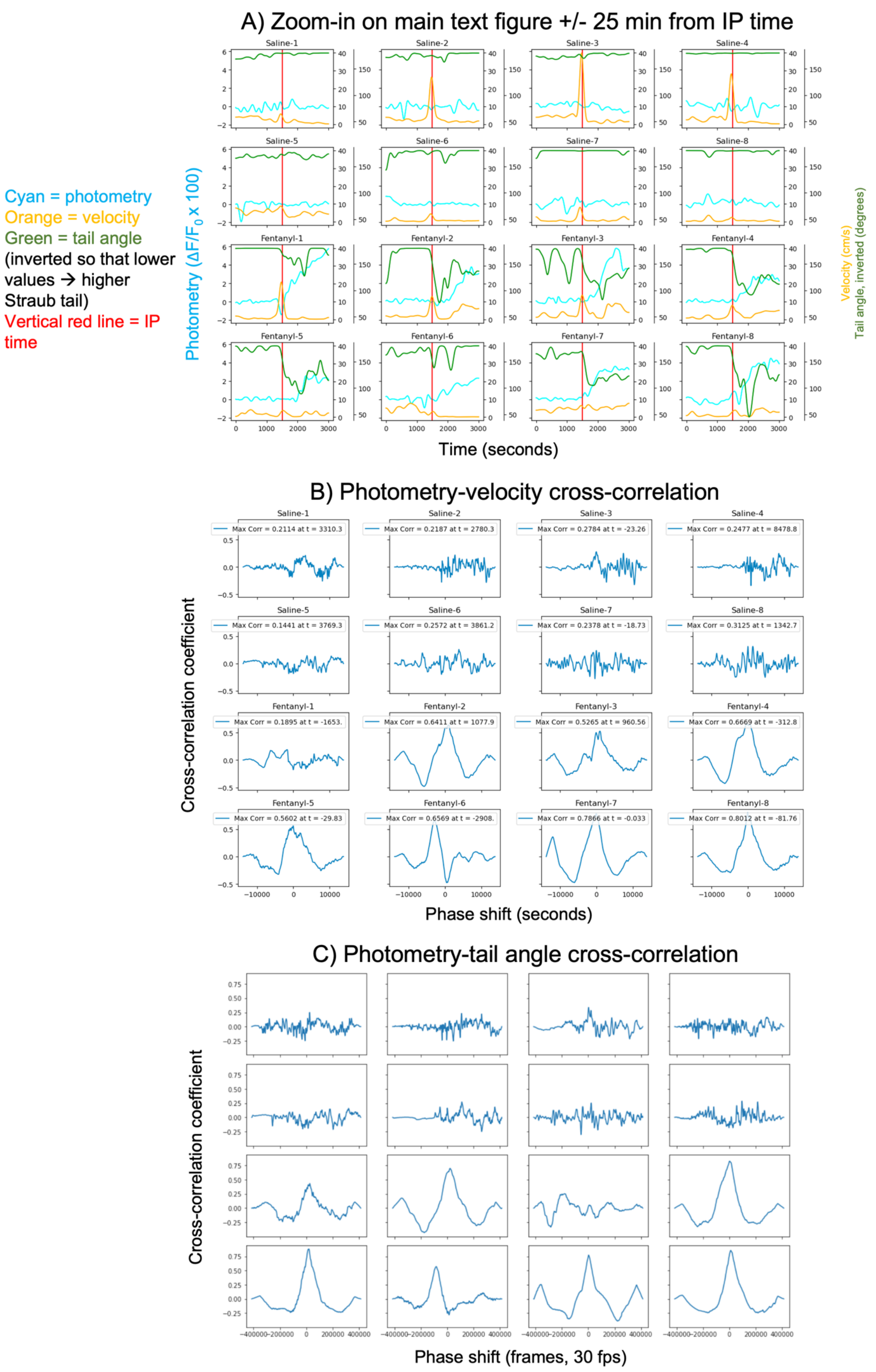
A) Zoom-in on main text figure +/- 25 min from IP time
Saline-1
Saline-2
Saline-3
Saline-4
Saline-5
Saline-6
Saline-7
Saline-8
Fentanyl-1
Fentanyl-2
Fentanyl-3
Fentanyl-4
Fentanyl-5
Fentanyl-6
Fentanyl-7
Fentanyl-8
Cyan = photometry
Orange = velocity
Green = tail angle
(inverted so that lower values → higher Straub tail)
Vertical red line = IP time
Photometry (ΔF/F0 x 100)
Velocity (cm/s)
Tail angle, inverted (degrees)
Time (seconds)
B) Photometry-velocity cross-correlation
Max Corr = 0.2114 at t = 3310.3
Max Corr = 0.2187 at t = 2780.3
Max Corr = 0.2784 at t = -23.26
Max Corr = 0.2477 at t = 8478.8
Max Corr = 0.1441 at t = 3769.3
Max Corr = 0.2572 at t = 3861.2
Max Corr = 0.2378 at t = -18.73
Max Corr = 0.3125 at t = 1342.7
Max Corr = 0.1895 at t = -1653.
Max Corr = 0.6411 at t = 1077.9
Max Corr = 0.5265 at t = 960.56
Max Corr = 0.6669 at t = -312.8
Max Corr = 0.5602 at t = -29.83
Max Corr = 0.6569 at t = -2908.
Max Corr = 0.7866 at t = -0.033
Max Corr = 0.8012 at t = -81.76
Cross-correlation coefficient
Phase shift (seconds)
C) Photometry-tail angle cross-correlation
Cross-correlation coefficient
Phase shift (frames, 30 fps)

**SI Figure 6 Related to Figure 5 & 6:**
(A) Zoom in on Figures 5 & 6 immediately before/after IP time with photometry (cyan), velocity (orange), and tail angle (dark green) traces overlaid. Straub tail metric is inverted with respect to the main text for clarity in the overlaid traces (depressed angle trace corresponds to a greater degree of Straub tail).
(B-C) Cross-correlations between photometry and behavior. X-axes are in video frames.
(B) Cross-correlation of the entire velocity vs. photometry waveforms.
(C) Cross-correlation of the entire tail angle vs. photometry waveforms.

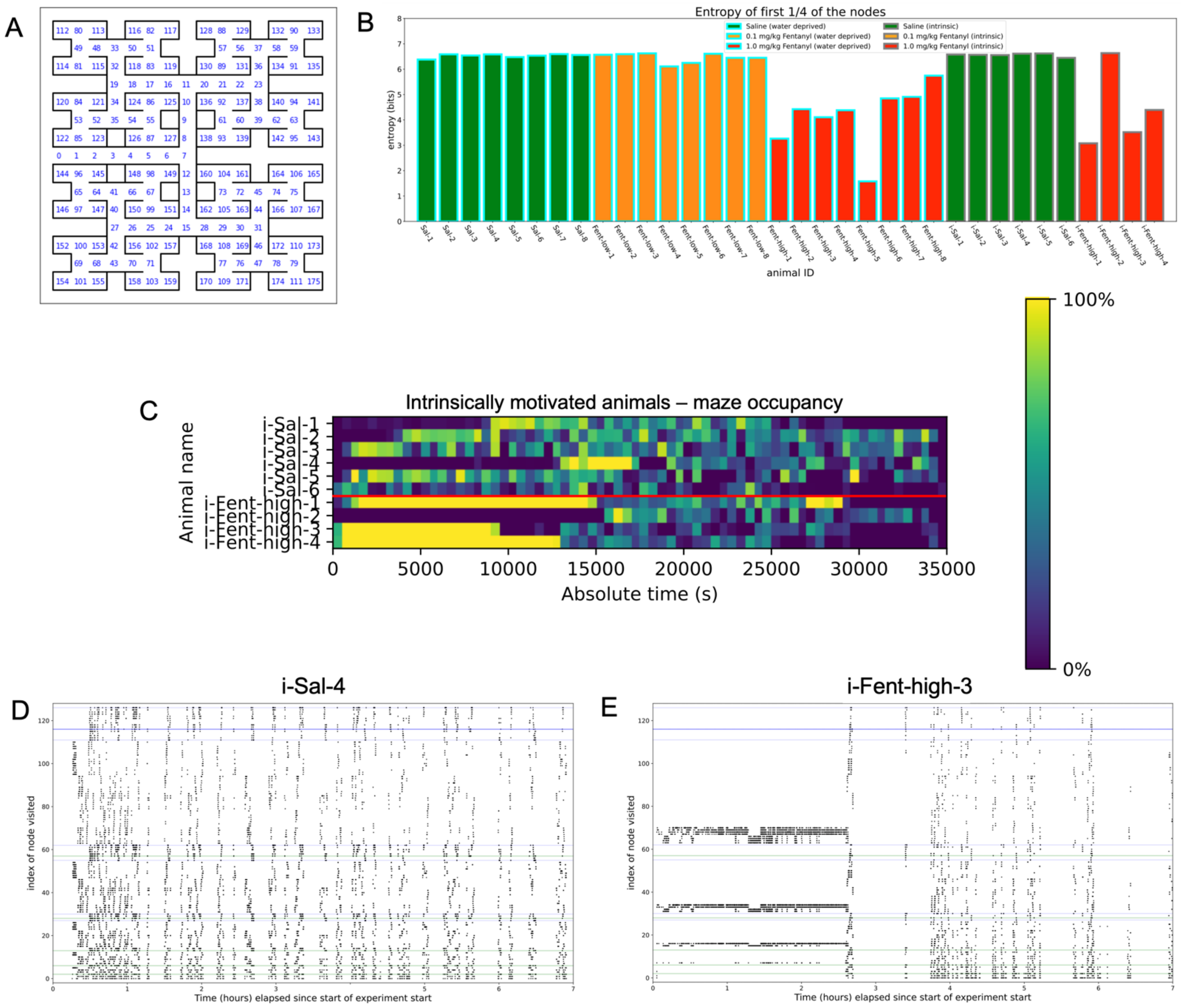

**SI Figure 7 related to Figure 7: additional entropy calculations and "intrinsic" negative control**
(A) Maze with node numbering used for Figure 7.
(B) Right: entropy of the maze navigation for the first ¼ nodes visited in the maze for each animal.
(C) "Intrinsic" negative control experiment – without water deprivation, animals receiving fentanyl still show repetitive navigation in the maze. Mice not deprived of water and given ad. lib. food and water in the home cage during the maze experiment. One group received saline IP (amA023, amA033, amB005, amB006, amB012, and amA034) and the other group received 1 mg/kg fentanyl IP (amA025, amB004, amB007, and amA020). Top: maze occupancy vs absolute time. Bottom: representative node navigation plots comparing an animal receiving saline IP (amB005) vs. an animal receiving 1 mg/kg fentanyl IP (amB004).

### STAR Methods:

#### KEY RESOURCES TABLE

| REAGENT or RESOURCE | SOURCE | IDENTIFIER |
| --- | --- | --- |
| Antibodies | | |
| Anti-GFP primary antibody | Abcam | Cat# ab13970 |
| Anti-tyrosine hydroxylase antibody | Millipore Sigma | Cat# AB1542 |
| Goat anti-chicken IgY secondary antibody, AlexaFluor488 | Abcam | Cat# ab150169 |
| Donkey anti-Sheep IgG secondary antibody, AlexaFluor568 | ThermoFisher Scientific | Cat# A-21099 |
| Bacterial and virus strains | | |
| AAV-hSyn-iFentanylSnFR2.0-cyto-WPRE | In house | N/A |
| NEB® Stable Competent *E. coli* (High Efficiency) | New England Biolabs | Cat# C3040H |
| *Escherichia coli* BL21(DE3) | Agilent Technologies, Santa Clara, CA | Cat# 200131 |
| One Shot™ TOP10 Electrocomp™ *E. coli* | ThermoFisher Scientific | Cat# C404052 |
| Biological samples | | |
| Hippocampal dissociated neuronal culture | C57BL/6NCrl mice; prepared in house | N/A |
| Acute brain slices | C57BL/6NCrl mice; prepared in house | N/A |
| Normal donkey serum | Abcam | Ab7475 |
| Normal goat serum | Abcam | Ab7481 |
| Human serum (pooled) | Lee Biosolutions | Cat# 991-58-P-RC |
| Human saliva (pooled) | Lee Biosolutions | Cat# 991-05-P |
| Human sweat | Gift from Wei Gao lab | N/A |
| Chemicals, peptides, and recombinant proteins | | |
| Alpha-endorphin | Avantor (VWR) | Cat# H-2695.0001BA<br>CAS: 59004-96-5 |
| Endomorphin-1 | Tocris | Cat# 1055<br>CAS: 189388-22-5 |
| Met-enkephalin acetate salt hydrate | Sigma Aldrich | Cat# M6638<br>CAS: 82362-17-2 |
| 7-hydroxymitragynine | Dalibor Sames lab | 174418-82-7 |
| Alfentanil hydrochloride | Cayman Chemical | Cat# 19292<br>CAS: 69049-06-5 |
| BMS986187 | Sigma Aldrich | Cat# SML0917<br>CAS: 313669-88-4 |
| BRL52537 hydrochloride | Tocris | Cat# 0699<br>CAS: 112282-24-3 |
| Butorphanol (+)-tartrate | Sigma Aldrich | Cat# B9156<br>CAS: 58786-99-5 |
| Codeine monohydrate | Sigma Aldrich | Cat# C5901<br>CAS: 6059-47-8 |
| Dextrorphan tartrate | Cayman Chemical | Cat# 15886<br>CAS: 143-98-6 |

| | | |
|---|---|---|
| Fentanyl citrate | Sigma Aldrich | Cat# F3886<br>CAS: 990-73-8 |
| GR89696 fumarate | Tocris | Cat# 1482<br>CAS: 126766-32-3 |
| Hydrocodone (+)-bitartrate salt | Sigma Aldrich | Cat# H4516<br>CAS: 143-71-5 |
| Hydromorphone hydrochloride | Sigma Aldrich | Cat# H5136<br>CAS: 71-68-1 |
| Levorphanol (+)-tartrate salt dihydrate | Sigma Aldrich | Cat# L5143<br>CAS: 5985-38-6 |
| Meperidine hydrochloride | Sigma Aldrich | Cat# M3142<br>CAS: 50-13-5 |
| Meptazinol hydrochloride | Tocris | Cat# 3636<br>CAS: 59263-76-2 |
| Mitragynine | Dalibor Sames lab | CAS: 4098-40-2 |
| ML138 | Sigma Aldrich | Cat# SML1140<br>CAS: 1355243-24-1 |
| Morphine sulfate pentahydrate | Sigma Aldrich | Cat# M8777<br>CAS: 6211-15-0 |
| Nalorphine hydrochloride | Sigma Aldrich | Cat# N-924<br>CAS: 57-29-4 |
| Naloxone hydrochloride dihydrate | Sigma Aldrich | Cat# N7758<br>CAS: 51481-60-8 |
| Naltrexone hydrochloride | Cayman Chemical | Cat# 15520<br>CAS: 16676-29-2 |
| Naltrindole hydrochloride | Cayman Chemical | Cat# 9000705<br>CAS: 111469-81-9 |
| (-)-nicotine tartrate | Fisher Scientific | Cat# BP25331<br>CAS: 65-31-6 |
| Norfentanyl | Cayman Chemical | Cat# 15899<br>CAS: 1609-66-1 |
| Oxycodone hydrochloride | Cayman Chemical | Cat# 26513<br>CAS: 124-90-3 |
| Remifentanil hydrochloride | Sigma Aldrich | Cat# R1908<br>CAS: 132539-07-2 |
| Salvinorin A | Cayman Chemical | Cat# 11487<br>CAS: 83729-01-5 |
| Sufentanil citrate | Sigma Aldrich | Cat# SML0535<br>CAS: 60561-17-3 |
| Tapentadol hydrochloride | Cayman Chemical | Cat# 9000620<br>CAS: 175591-09-0 |
| Tramadol hydrochloride | Sigma Aldrich | Cat# 1672600<br>CAS: 36282-47-0 |
| Acetylcholine chloride | Sigma Aldrich | Cat# A2661<br>CAS: 60-31-1 |
| Choline chloride | Sigma Aldrich | Cat# C7527 |
| Dopamine hydrochloride | Sigma Aldrich | Cat# H-8502 |
| γ-aminobutyric acid (GABA) | Sigma Aldrich | Cat# A2129 |
| Gluatmate | Sigma Aldrich | Cat# G1251 |
| Glycine | Sigma Aldrich | Cat# G7126 |
| Histamine dihydrochloride | Sigma Aldrich | Cat# H7250 |
| (-)-norepinephrine bitartrate | Cayman Chemical | Cat# 16673 |
| Serotonin creatine sulfate monohydrate | Fluka | Cat# 85030 |
| Lysogeny broth - BD Difco™ LB Broth | Lennox | Cat# 240230 |
| LB with Agar - BD Bacto™ Agar | Sigma Aldrich | Cat# 214010 |
| Ampicillin sodium salt | Sigma Aldrich | Cat# A0166-5G |
| Hanks' Balanced Salt Solution (HBSS) | Gibco | Cat# 14025-092 |

| | | |
|---|---|---|
| PBS (10x), pH 7.4 | Thermo Fisher Scientific | Cat# 70011044 |
| DPBS 1x (no calcium, no magnesium) | Gibco | Cat# 14190-250 |
| DMEM (high glucose, GlutaMAX supplement, pyruvate) | Gibco | Cat# 10569-044 |
| Fetal bovine serum | Gibco | Cat #26140079 |
| Penicillin–streptomycin (5,000 U/ml) | Gibco | Cat# 15070-063 |
| Trypsin-EDTA (0.25%), phenol red | Gibco | Cat# 25200072 |
| Lipofectamine 3000 Transfection Reagent | Thermo Fisher Scientific | Cat# L3000015 |
| Opti-MEM™ | Gibco | 31985070 |
| 35 mm dishes with optical element coated with poly-D-lysine | MatTek | P35GC-1.5-10-C |
| poly-L-ornithine | Sigma | P4957-50ML |
| laminin | Sigma | L2020-1MG |
| WFI for cell culture | Gibco | Cat# A1287304 |
| Polyethylenimine, linear, MW 25,000 | Polysciences | Cat# 23966-100 |
| OptiPrep - iodixanol gradient | Cosmo Bio USA | Cat# AXS-1114542-5 |
| Polyethylene glycol (MW 8,000) | Sigma Alrich | Cat# 89510 |
| Salt-active nuclease (25 U/μl) | ArcticZymes | Cat# 70910-202 |
| Pluronic F-68 | Gibco | Cat# 24040-032 |
| SmaI | New England Biolabs | Cat# R0141 |
| ScaI-HF | New England Biolabs | Cat# R3122S |
| Proteinase K | Roche Diagnostics | Cat# 03115828001 |
| DNase I recombinant (RNase-free; 10 U/μl) | Roche Diagnostics | Cat# 4716728001 |
| N-lauroylsarcosine sodium salt | Sigma Aldrich | Cat# . L9150 |
| Isoflurane, USP | Piramal Critical Care | Cat# 66794-017-25 |
| VECTASHIELD Vibrance® Antifade Mounting Medium with DAPI | Vector Laboratories | H-1800-2 |
| 0.9% sodium chloride injection, USP | Hospira | Cat# NDC 0409-4888-02 |
| Critical commercial assays | | |
| Phusion High-Fidelity PCR Kit | New England Biolabs | E0553L |
| QIAprep Spin Miniprep Kit | QIAGEN | 27104 |
| EndoFree Plasmid Maxi Kit | QIAGEN | 12362 |
| QIAquick Gel Extraction Kit | QIAGEN | 28704 |
| QIAquick PCR Purification Kit | QIAGEN | 28104 |
| Gibson Assembly® Master Mix | New England Biolabs | E2611 |
| Metabond® Quick Adhesive Cement System | Parkell | S380 |
| Qubit dsDNA HS Assay Kit | Invitrogen | Cat# Q32854 |
| SYBR Green master mix | Roche Diagnostics | Cat# 04913850001 |
| Deposited data | | |
| Raw and analyzed data (dose response, photometry, videography, and machine vision annotation). | This paper | |
| pHHM-iTapentadolSnFR | This paper | |
| pHHM-iLevorphanolSnFR | This paper | |
| pHHM-iFentanylSnFR1.0 | This paper | |
| pHHM-iFentanylSnFR2.0 | This paper | |
| pMinDisplay-iFentanylSnFR1.0-PM | This paper | |
| pMinDisplay-iFentanylSnFR1.0-ER | This paper | |
| pMinDisplay-iFentanylSnFR2.0-PM | This paper | |
| pMinDisplay-iFentanylSnFR2.0-ER | This paper | |
| pAAV-iFentanylSnFR2.0-cyto-WPRE | This paper | |
| iNicSnFR3a crystal structure | PDB: 7S7V | |

| | | |
|---|---|---|
| Experimental models: Cell lines | | |
| HeLa | ATCC | CCL-2; RRID:CVCL_0030 |
| HEK293T | ATCC | CRL-3216 |
| Experimental models: Organisms/strains | | |
| *Mus musculus* C57BL/6NCrl, 8-12 weeks old | Charles River | strain code 027 |
| Oligonucleotides | | |
| hSyn-forward | CCCGCAAACTCCCCTTC | |
| WPRE-forward | GGCTGTTGGGCACTGACAAT | |
| WPRE-reverse | CCGAAGGGACGTAGCAGAAG | |
| Recombinant DNA | | |
| pHHM.X513-iNicSnFR3b-(V7) | Addgene | #124881 |
| pMinDis.X513-iNicSnFR3a-(CC93)-ER | Addgene | #125121 |
| pMinDis.X513-iNicSnFR3a-(CC93)-PM | Addgene | #125122 |
| pAAV-hSyn-iGluSnFR-WPRE-SV40 | Addgene | #98929 |
| pHelper | Agilent | Cat: 240071 |
| AAV9 capsid | AddGene | 112865 |
| Software and algorithms | | |
| Python, version 3.8.5 | Python Software Foundation | https://www.python.org |
| Opioid chemoinformatics | This work | TBD |
| Behavior analysis pipeline | This work | TBD |
| Origin Pro, version 9.1 | OriginLab | |
| AutoDock Vina, version 1.2.0 | The Scripps Research Institute's Molecular Graphics Laboratory | |
| AutoDockTools, version 1.5.6 | The Scripps Research Institute's Molecular Graphics Laboratory | |
| ChimeraX, version 1.2.5 | Resource for Biocomputing, Visualization, and Informatics at the University of California, San Francisco | |
| PyMol, version 2.5.0 | Schrödinger LLC | |
| ColabFold (MMseqs2 and AlphaFold2) | | https://github.com/sokrypton/ColabFold |
| Tecan plate reader software, version 3.1 | Tecan | |
| iQ3 (image acquisition) | Andor | N/A |
| ZEN Microscopy | Zeiss | N/A |
| DeepLabCut | | |
| Labyrinth maze analysis Python code | Markus Meister Lab | https://github.com/markusmeister/Rosenberg-2021-Repository |

| | | |
|---|---|---|
| NanoAnalyze | TA Instruments | https://www.tainstruments.com/sw/nano_analyze.html |
| KaleidaGraph | Synergy | RRID:SCR_014980 |
| NEBuilder | New England Biolabs | https://nebuilder.neb.com/ |
| MATLAB R2016a/2017b | MathWorks | N/A |
| Other | | |
| Narcan (Naloxone hydrochloride nasal spray) | Adapt Pharma, Inc. | Cat# NDC 69547-353-02 |
| Analytical balance | Mettler & Toledo | XPR204 |
| 2.2 mL 96-well storage plates | Thermo Scientific | Cat# AB0932 |
| AeraSeal film | Sigma Aldrich | Cat# A9224 |
| Sonicator (Sonifier SFX550) | Branson | Cat# 101-063-969 |
| Spark 10M multimode plate reader | Tecan | Cat# 30086376 |
| epMotion 5075t liquid handling robot | Eppendorf | Cat# 5075006022 |
| Spectrophotometer | Thero Scientific | Nanodrop 1000 |
| ÄKTA Start protein purification system | Cytiva | Cat# 29023051 |
| HiTrapTM 5ml IMAC FF Nickel NTA column | Cytiva | Cat# 17-0921-04 |
| 50 mL Amicon Ultra centrifugal filter tubes (30 kDa cutoff) | Sigma Aldrich | Cat# UFC903096 |
| Protein LoBind tubes (2.0 mL) | Eppendorf | Cat# 022431102 |
| Sterile BioStor™ vials with screw cap | National Scientific Supple Co, Inc. | Cat# BC16NA-PS |
| Isothermal titration calorimeter | TA Instruments | Affinity ITC |
| Stopped-flow fluorimeter | Applied Photonics | SX20 |
| Inverted widefield fluorescence microscope | Olympus | IX-81 |
| Back illuminated electron multiplier CCD camera | Andor Technology | iXon DU-897 |
| 470 nm LED | Led Engin | Cat# LZI-10DB00 |
| 40 nm band-pass filter centered on 470 nm | Chroma Technology | Cat# ET 470/40X |
| Automated perfusion controller | AutoMate Scientific | ValveBank8 |
| Micro-incubator with perfusion tubes | Warner Instruments | Cat# DH-40i |
| OptiSeal tubes | Beckman Coulter | Cat# 361625 |
| Thermal cycler | BioRad | C1000 Touch |
| Real-time detection system for qPCR | BioRad | CFX96 |
| Vibrating microtome (Compresstome®) | Precisionary Instruments | Cat# VF-310-0Z |
| iSpacer® | SunJin Lab Co. | Cat# IS203 |
| Slice anchor | Warner Instruments | SHD-26GH/15 |
| Upright microscope | Olympus | BX50WI |
| Transmitted light source (400-700 nm) | Sutter Instruments | TLED+ |
| Digital CCD camera | Hamamatsu | ORCA-03G |
| Zeiss Axio Observer 7 with Definite Focus 2 with Airyscan 2 Module | Zeiss | LSM980 |
| Small animal stereotactic surgical apparatus | Stoetling | ??? |
| Micro4 controller | World Precision Instruments | UMC4 |
| Nanoliter injector | World Precision Instruments | ??? |
| 33g blunt NanoFil needle | World Precision Instruments | Cat# NF33BL |
| Sub-microliter injection system syringe (10 $\mu$L) | World Precision Instruments | Cat# NANOFIL |
| Fiber Optic Cannula, Ø2.5 mm Ceramic Ferrule, Ø400 $\mu$m Core, 0.39 NA, L=10 mm | ThorLabs | CFMC14L10 |
| Ruby DualScribe Fiber Optic Scribe | ThorLabs | S90R |

| | | |
|---|---|---|
| Fiber Inspection Scope | ThorLabs | FS201 |
| Vetbond Tissue Adhesive | 3M | 1469SB |
| Denture Repair Powder | Lang Dental Manufacturing | Jet Set-4 |
| Jet Liquid | Lang Dental Manufacturing | Jet Liquid |
| [Oka photometry setup] | | |
| 490 nm LED (photometry) | Thorlabs | Cat# M490F1 |
| 405 nm LED (photometry) | Thorlabs | Cat# M405F1 |
| Photoreceiver | Newport | Cat# 2151 |
| Real-time digital signal processor | Tucker-David Technologies | Cat# RP2.1 |
| Video camera – 1080p | Logitech | c920s HD Pro |
| White light bar (3000-5000K) | Defiant | Cat# YT-8001C |
| Bpod State Machine r1 (precision animal behavior measurement platform) | Sanworks | Cat# 1027 |
| Mouse port assembly (IR beam break, valve, and mount) for liquid reward delivery | Sanworks | Cat# 1009 |
| IR illuminator (850 nm, 12 LED, wide angle) | Univivi | Cat# 4331910725 |
| 0.3 mL syringes with 29G needle | Becton Dickinson | Cat# 324702 |
| Graphical processing unit | NVIDIA | GeForce GTX 1080 Ti |

#### RESOURCE AVAILABILITY

##### Lead contact

#### EXPERIMENTAL MODEL AND SUBJECT DETAILS

##### Mice

All animal care and experimental procedures were carried out in accordance with the National Institutes of Health (NIH) Guide for the Care and Use of Laboratory Animals and approved by the Institute Animal Care and Use Committee and the Institute Biosafety Committee at the California Institute of Technology. Both males and females were used in equal numbers in each experiment. The animals were housed in a temperature- and humidity-controlled facility with a 13:11 h light:dark cycle with food and water available *ad libitum*. The animals were group-housed when possible. Following surgery, subjects were singly housed for two weeks prior to experiments and monitored daily for a full recovery. Mice in the experimental arm of the maze foraging experiment were deprived of water for 21 h before the experiment.

#### METHOD DETAILS

##### Cloning and DNA Preparation

###### *Cloning*

Gibson assembly was used to construct several biosensor-vector combinations. We used NEBuilder (https://nebuilder.neb.com/) to design primers to generate DNA fragments with the requisite overlaps. These fragments were combined using Gibson assembly reactions (Gibson Assembly® Master Mix, NEB) according to the manufacturer's protocol. Each type of vector, bacterial, mammalian, and viral, served different experiments and dictated additional steps:

*Bacterial expression vector*: We used our previously reported vector pHHM.X513-iNicSnFR3b-(V7) (Addgene #124881) and replaced iNicSnFR3a with the new gene of interest. Directed evolution experiments mutated the resulting plasmids directly.

*Mammalian expression vectors*: we used our previously reported pMinDis.X513-iNicSnFR3a-(CC93)-ER (Addgene #125121) and pMinDis.X513-iNicSnFR3a-(CC93)-PM (Addgene #125122), targeting the endoplasmic reticulum and plasma membrane, respectively. We replaced the gene for iNicSnFR3a with the new biosensor gene of interest.

*pAAV construction for AAV packaging*: pAAV-hSyn-iGluSnFR-WPRE-SV40 (AddGene #98929) was used as the backbone for the viral construct. The PDGFR-targeting sequence was excluded in the final construct. Without any targeting sequences, biosensors are directed to the cytoplasm of neurons. PCR reactions involving the vector backbone required 3% DMSO due to the secondary structure of the inverted terminal repeat (ITR) regions. SmaI digest was used to confirm the integrity of the ITR regions. The resulting plasmid, pAAV-hSyn-iFentanylSnFR2.0-cyto-WPRE, was used for AAV production.

###### *DNA preparation and purification*

Plasmids used for bacterial expression were transformed into DH5α cells and cultured on ampicillin selection plates for 16-20 h at 37 °C. A 5 mL LB culture with ampicillin was inoculated with one colony and incubated while shaking at 250 rpm at 37 °C for 16-18 h. The DNA was purified using the QIAPrep Spin Miniprep Kit (Qiagen) according to the manufacturer's protocol. The resulting DNA was submitted for Sanger sequencing to verify its identity (Laragen Inc., Culver City, CA).

pAAV plasmids used for AAV production were transformed into NEB Stable cells using the NEB Stable media and cultured on ampicillin selection plates at 30 °C for 30 h. All culturing tubes and flasks were pyrogen-free and specified for AAV production. A 5 mL primary culture using Plasmid+® media (Thomson Instrument Company) and ampicillin was inoculated with a single colony and incubated while shaking at 250 rpm at 30 °C for 6-8 h. The primary culture was diluted 1:200 into a secondary culture of 100 mL Plasmid+® media with ampicillin and incubated while shaking at 250 rpm at 30 °C for 16-20 h. The DNA was purified using EndoFree Plasmid Maxi Kit (Qiagen) according to the manufacturer's protocol. SmaI digest was used to confirm the integrity of the inverted terminal repeat (ITR) regions. The region between the ITRs was sequenced using custom hSyn-forward and WPRE-reverse primers (Laragen Inc., Culver City, CA). The DNA concentration and purity were verified using a Nanodrop.

#### Drug solution preparation

##### *Safety for handling fentanyl and its analogs*

During the procedure, the experimenter wore a disposable lab coat, sleeve covers, a safety mask, and eye protection. Two observers, trained in administering Narcan and in emergency response procedures related to opioid exposure, stood by for the duration of the procedure. The powder was weighed, transferred to a container, and dissolved in an aqueous solution. The bench space was wiped down with water and 70% ethanol solution. The plastic waste was triple-rinsed, and all waste was disposed of using standard chemical hazard procedures.

##### *Stock solution preparation*

Controlled substances were procured under a Schedule II DEA license (PI: Henry Lester) and stored in lockboxes at room temperature, 4 °C, or –20 °C, according to the manufacturer's suggested conditions. An analytical balance with 0.0001 g precision was used to weigh compounds. Stock solutions were prepared in 3x PBS pH 7.0 or sterile saline for *in vitro* and *in vivo* experiments, respectively. Drug solutions were sterile-filtered using a 0.2- µM syringe filter before injections in mice. Stock solutions were serially diluted and stored in deep well plates for dose-response experiments. All solutions and plates were stored at –20 °C.

#### Protein Purification and Lyophilization

##### *Protein purification*

A bacterial expression vector bearing the biosensor gene of interest was used to transform chemically competent BL21 DE3 gold cells (Agilent Technologies). Bacteria were plated on LB agar selection plates (100 mg/L ampicillin). A single colony was used to inoculate 200 mL of autoinduction media (Studier 2005) with 100 mg/L ampicillin. The culture was incubated at 30 °C while shaking at 250 rpm for 28-30 h. The resuspended cell pellet was sonicated on ice three times, 30 sec each, with 2 min recovery periods in between. The cell debris was pelleted by centrifugation. The supernatant was filtered through a 0.2 *µ*m filter and loaded onto an *Ä*KTA Start FPLC equipped with a 5 mL Ni-NTA column. The protein was eluted using a 10-200 mM imidazole gradient in 1x PBS, pH 7.4. The fractions were analyzed by SDS-PAGE gel for the expected mass and fraction purity. Pure fractions were pooled, concentrated, and buffer exchanged into 3x PBS pH 7.0 using a spin column with a 30 kDa cutoff (Amicon). Protein concentration was determined by measuring the sample's absorbance at 280 nm. Purified protein was stored at 4 °C.

##### *Lyophilization and simulated field test*

100 µL samples of the purified fentanyl biosensor were flash-frozen in liquid nitrogen and lyophilized. The biosensor powder was stored in the dark to prevent the photobleaching of GFP. The tube containing the powder was punctured so that the samples could be exposed to room temperature and humidity. After three weeks, the powder was redissolved into 100 µL molecular biology grade deionized water, and the concentration was verified unchanged by this process. A control sample from the same batch of purified protein was flash-frozen and stored at – 80 °C during the same period. The lyophilized and control biosensor samples were then tested using the general dose-response method.

#### General methods of dose-response measurements

##### *General Tecan plate reader method*

Plates were read using a Tecan Spark 10M with 485 nm excitation (20 nm bandwidth) and 535 nm emission (25 nm bandwidth) wavelengths in top read mode and manual gain set to "60" to measure GFP fluorescence. Samples were mixed by pipette in the plate and then inserted into the instrument, where it is shaken for 10 seconds using a double orbital pattern prior to recording from the designated wells. All dose-response experiments were carried out at room temperature.

##### *Dose-responses in PBS*

Purified biosensor protein and drug solution plates were mixed using a robotic liquid handler (epMotion, Eppendorf). 11 µL of a drug solution from the 10x concentration stock plate was mixed into 100 µL biosensor solution in 3x PBS pH 7.0 in triplicate for each dose. The concentrations of biosensor and drug were chosen to observe interactions surpassing the ligand depletion regime. The final [biosensor] = 100 nM in each well for experiments involving purified protein. One set of three wells was reserved for buffer control (zero [drug] to determine $F_0$).

*Dose responses in biofluids*

Biofluids were frozen and thawed once to aliquot prior to the experiment and were not filtered, pH-adjusted, or otherwise modified, thereby maintaining their composition. A series of biosensor/drug solutions were prepared with 2x the final target concentration in a final volume of 50 $\mu$L. These solutions were manually pipetted into a Costar flat black 96 well plate. The biofluid was pipetted and mixed thoroughly in each well to yield the final drug-spiked biofluid-biosensor solution. The fluorescence was read using the general Tecan plate reader method.

#### Directed Evolution

*Mutant library generation by site saturation mutagenesis (SSM)*

Residues for site saturation mutagenesis were chosen based on previously reported crystal structures and mutagenesis data. Generally, sites in the linker region and hinge were chosen to improve dynamic range, and sites in the binding pocket and second shell were chosen to improve affinity for the target opioid. DNA libraries were constructed using the "22-codon method," where three sets of primers bearing 'NDT', 'VHG', and 'TGG' encode for 12, 9, and 1 codons, respectively, at the site of interest (Kille 2013). The 22-codon method covers all 20 amino acids with near-equal selection probability while excluding all stop codons. A PCR reaction using Phusion polymerase, an equimolar solution of each primer, and the biosensor parent plasmid as the template DNA generated the "SSM library". The PCR product was purified and treated with a Dpn1 digest to remove all template DNA. The DNA libraries were transformed into electrochemically competent TOP10 cells and plated on LB agar ampicillin selection plates. Typically, we found ~300 colonies on a plate, ensuring sufficient sampling of the DNA library. Five colonies were selected at random to inoculate miniprep cultures (5 mL LB with ampicillin). The DNA was isolated and sequenced to verify codon randomization from the 22-codon method. The remainder colonies on the selection plate were resuspended in LB and centrifuged. The DNA was extracted using a miniprep kit to yield an amplified SSM DNA library.

*Bacterial lysate screening of mutant libraries*

Separate aliquots of BL21 DE3 cells were transformed with 300 ng of the SSM library and the parent biosensor plasmid and cultured on LB agar ampicillin (100 mg/L) selection plates. Autoinduction medium was prepared according to the Studier method (Studier 2005), and 800 $\mu$L was pipetted into each well in a 96-deep well plate. 92 wells were inoculated with a colony randomly picked from the SSM library plate, three with colonies with the parent biosensor (positive control), and one with no inoculation (negative control). The plate was sealed with a 0.2 $\mu$m breathable mesh (AeraSeal, Sigma Aldrich) and incubated while shaking at 250 rpm at 30 °C for 28 h. After culturing, 100 µL from each well was transferred to the corresponding wells in a 96-well Costar flat black plate and stored at -80 °C to create a replica plate. The remainder culture in deep well plate was centrifuged to yield a yellow-green pellet. The supernatant was removed, and the pellet was washed once with 1x PBS pH 7.0 to remove residual media. The pellet was then resuspended in 3x PBS pH 7.0, flash-frozen in liquid nitrogen, and thawed to lyse the cells. The plate was centrifuged again, yielding solubilized mutant biosensors in the lysate.

The mutants were then screened in parallel using positive and negative screens (if applicable) against the opioid of interest and acetylcholine, respectively. The concentrations of the ligands used in the screens were their $EC_{50}$ in activating the parent biosensor. 100 µL of the lysate from each well was transferred to each Costar flat black plate. The plate reader was programmed to read GFP fluorescence, add 11 µL of a 10x stock of the ligand to each well, shake in a double orbital pattern to mix, and then read GFP fluorescence again. These data provided the $F_0$ and $\Delta F$ values to determine the strength of the response in each well. The DNA of mutants that showed considerable improvement was isolated by culturing the BL21 DE3 from the replica plate, extracting the DNA using a miniprep kit, and sequencing to determine the winning residue identity.

*Full dose responses in lysate for winning mutants*

To compare the winning mutations from the plate screen, the top several constructs were compared in full dose responses before selecting one for further directed evolution. BL21 DE3 cells were transformed with the DNA of the winning mutants and plated on selection plates with ampicillin. A 10 mL culture with autoinduction media and ampicillin was inoculated for each winner and incubated at 30 °C while shaking (250 rpm) for 28 h. The culture was centrifuged to yield a yellow-green pellet. The pellet was resuspended in 8 mL of 3x PBS pH 7.0, flash-frozen in liquid nitrogen, and thawed to lyse the cells. The lysate was centrifuged, and the supernatant was pipetted to yield the solubilized biosensor. A serial dilution of the lysate was performed, and fluorescence was measured on the plate reader to determine the appropriate dilution that provided a baseline fluorescence of ~1/20 of the detector's dynamic range under the given settings. Each mutant was characterized using the general dose-response method against the opioid of interest and acetylcholine (if applicable). The mutant displaying the greatest improvement was taken forward for the next iteration of site saturation mutagenesis at another set of residues.

#### Biophysical Characterization

##### *Isothermal titration calorimetry*

The Affinity ITC (TA Instruments) equipped with a 190 µL cell was used for all isothermal titration calorimetry experiments. Biosensors were purified and buffer-exchanged into 3x PBS pH 7.0. Biosensor concentrations in the range of 20-50 µM were sufficient to produce a large enough dynamic range between the first injection and final heats. The same batch of buffer was used to dilute the protein and prepare drug solutions used in the ITC experiment. The drug solution was prepared for a final concentration ten times that of the biosensor's final concentration in the ITC cell. All solutions were degassed before the experiment. The biosensor solution was added to the cell, and the drug solution to the syringe (titrant). 2-2.5 µL injections of the titrant were injected at 300 s intervals 20 times, spanning 0 to 2.0-2.5 mol equivalents of the opioid to the biosensor. NanoAnalyze software (TA Instruments) was used to process and fit the data. The integrated heats were fit with the "independent" plus "constant" models to account for the drug-biosensor binding interaction and the drug solvation energy, respectively. The resulting fit from this model determined enthalpy, entropy, binding affinity, and stoichiometry for each biosensor-opioid pair.

##### *Stopped-flow kinetics experimentation and data analysis*

Stopped-flow kinetics were measured using a stopped-flow fluorimeter equipped with a 490 nm excitation LED and 510 nm long-pass filter (Applied Photophysics SX20) at room temperature (22 °C). Equal volumes of 0.2 µM biosensor and varying drug concentrations were mixed (5 replicates). The first 3 ms were not included in the final analysis and fitting to isolate mixing artifacts and instrument dead time. Data were plotted, and time courses were fitted using Kaleidagraph (version 4.4). $k_{obs}$ were plotted as a function of [ligand], and the linear regime was fitted; the slope reporting $k_1$ and the y-intercept reporting $k_{-1}$. When the time course did not fit well to a single exponential component, it was fitted to the sum of two exponentials, and the faster phase ($k_{obs1}$) was treated as above to determine $k_1$ and $k_{-1}$.

#### Cell Line Imaging

##### *HeLa cell tissue culture and transfection*

HeLa cells were procured from ATCC and cultured according to their suggested protocol: cultured in 10% EMEM in FBS supplemented with penicillin/streptomycin media at 5% $CO_2$ and 37 °C. Medium changes were performed when flasks reached ~80% confluence (~48 h). The cells were thawed into a flask and passaged at least twice before any experiments. For imaging experiments, 100,000 HeLa cells were plated onto each 35 mm dish with a built-in 14 mm coverslip (MatTek) and incubated for 24 h. The cells were then transfected with Lipofectamine 3000 and either 100 ng of iFentanylSnFR1.0_ER or 500 ng of iFentanylSnFR1.0_PM in OptiMEM. The cells were incubated in the transfection medium for 18 h and then incubated in the standard growth medium for 24 h before the imaging experiment.

##### *Imaging under fentanyl bath perfusion*

The cells were then imaged under widefield epifluorescence (40x, 1.0 NA) on an inverted microscope (IX-81, Olympus) equipped with a camera (Andor) and a gravity-fed eight-channel perfusion system (Automate Scientific). Imaging was performed in widefield epifluorescence mode and recorded at 4 Hz using Andor's iQ3 software. A programmable perfusion controller (Automate Scientific) was used to set the sequence and duration of perfusion. The chamber has inlet and outlet tubes spaced ~3 mm apart and are shaped to provide laminar flow across the cells in between. The solution change time is ~3 s. Generally, a 1 min HBSS control followed by a 1 min washout was applied to determine any artifacts from changes in pH due to bicarbonate exchange with the environment. Each drug dose was applied for 2 min and washed out for 2 min. The GFP response to the artifactual pH transient was subtracted to provide the true response to the drug perfusion as we have previously described (Shivange 2019).

#### Virus Production and Titering.

##### *Virus generation and purification*

We followed the protocol reported by Challis et al. (2019) the produce the virus used in this work. All the reagents listed in the protocol were used as stated, and only the vector and gene of interest were changed to pAAV-hSyn-iFentanylSnFR2.0-cyto-WPRE. HEK293T cells were transfected with the pAAV and two other plasmids, pHelper and AAV9 capsid, to assemble the viral particles. The media was changed at one and three days after transfection. The media from the second change was saved, and the cells and media were harvested after five days and combined with the saved media. The entire suspension was centrifuged to separate intact cells from the media. A polyethylene glycol (PEG) solution was added to the media, incubated for 2 h, and centrifuged to separate viral particles into the PEG pellet. The cell pellet was resuspended in a buffer with salt active nuclease (SAN) and incubated for 1 hour. The PEG pellet was resuspended and added to the cell suspension for further SAN digestion.

An OptiSeal tube was prepared with an iodixanol gradient layered with 15%, 25%, 40%, and 60% weight/weight iodixanol in DPBS. The virus was loaded into the OptiSeal tube and purified by ultracentrifugation. The tube was removed from the rotor, and the layer bearing the virus was removed by a needle and syringe puncturing the tube. The virus was buffer exchanged to remove iodixanol and equilibrate into sterile DPBS with Pluronic F68 surfactant. The virus was stored in a sterile, low protein-binding screw cap vial at 4 °C. A typical yield from 10 x 15 cm culture dishes was a purified virus sample of ~0.6 mL at a concentration of ~$1.0 \times 10^{13}$ vg/mL.

##### *Titering by qPCR*

The titer was quantified before each batch of surgical injections. Briefly, a sample of the virus was treated with DNAse followed by Proteinase K to isolate only the packaged DNA. A serial dilution of the packaged DNA was mixed with a SYBR green qPCR master mix (Roche). qPCR reaction was conducted on a thermocycler (C1000 Touch, BioRad) with a real-time detection system (CFX96, BioRad) using the protocol suggested by Challis et al.:

Step 1: 95 °C, 10 min
Step 2: 95 °C, 15 s
Step 3: 60 °C, 20 s
Step 4: 60 °C, 40 s
Repeat steps 2–4 40×.

#### Dissociated hippocampal neuronal culturing, transduction, and imaging

24 h prior to the dissociation, MatTek dishes with a 10 mm glass coverslip coated with poly-D-lysine were further coated with poly-L-ornithine and laminin. The next day, a pregnant mouse (C57BL/6NCrl, Charles River) was euthanized at embryonic day 16, and the uterine sac was removed. Each embryo was decapitated and then dissected. Hippocampi from several embryos were pooled and digested with 15 units of papain for 15 min at 37 °C and then treated with DNase. The dissociated cells were triturated in HBSS with 5% donor equine serum and centrifuged through a layer of 4% BSA in HBSS. The cells were plated on the treated dishes at a density of 90,000 cells/dish in 130 µL of plating medium deposited on the coverslip. After 1 h, 3 mL of complete culture medium was added to fill the dish. Medium changes were performed twice a week. Four days after plating, the neurons were transduced with virus at a multiplicity of infection of 50,000. After 2 weeks, the neurons stably expressed the biosensor, and the dishes were imaged as in the HeLa cell experiments.

#### Slice Experiments

##### *Surgical procedure*

Mice aged 8-12 weeks underwent anesthesia induction with 5% isoflurane in oxygen and were maintained under anesthesia at 1-2% isoflurane. Mice were kept on a heating pad during the surgery to maintain body temperature. The mice were mounted on a small animal stereotaxic apparatus (Stoelting) equipped with ear bars. The skull was leveled to place lambda and bregma on the same z-axis. The coordinates for the VTA used in this study were AP -3.30; ML +0.40; DV −4.30. A drill bit attached to the stereotactic rig was calibrated based on lambda and bregma and then used to drill a hole through the skull. Saline was used to flush the opening. A sub-microliter injection system with a 33 g needle (World Precision Instruments) attached to a micrometer controller (World Precision Instruments) was backfilled with oil, loaded with the virus (AAV9-hSyn-iFentanylSnFR2.0-cyto-WPRE), and then lowered into position over the course of ~7 min. 250 nL of virus (2.5 x $10^9$ vg) was injected at 100 nL/min, regulated by a MicroPump 4. We waited for 5 min to allow the virus to diffuse in the tissue before removing the needle. After the surgery, mice were placed in a new cage with a heating pad underneath half of the cage and observed until they recovered from the anesthesia. Thereafter, the mice were checked daily for a routine recovery. The mice were allowed > 2 weeks of recovery before the acute slice experiments.

*Acute brain slicing*

We followed the method of Ting et al. 2018 "Preparation of Acute Brain Slices Using an Optimized *N*-Methyl-D-glutamine Protective Recovery Method," and the manufacturer's protocols for operating the Compresstome (Precisionary Instruments). Solutions were prepared, their osmolarity adjusted according to Ting 2018, and temperature equilibrated prior to handling the animal. Animals were anesthetized, and once non-responsive, transcardiac perfusion was performed. The animal was decapitated and then the brain was removed into cold NMDG-HEPES aCSF (in mM: NMDG, 92; HCl, 92; KCl, 2.5; $NaH_2PO_4$, 1.2; $NaHCO_3$, 30; HEPES, 20; glucose, 25; sodium ascorbate, 5; thiourea, 2; sodium pyruvate, 3; $MgSO_4$, 10; $CaCl_2$, 0.5). The brain was sectioned and mounted on a block. Molten agar was poured to submerse the brain and then rapidly cooled over ~few seconds. The brain was cut into 200 $\mu$m coronal slices while the block was bathed in cooled NMDG-HEPES aCSF solution. Beginning with the cutting step through the end of the experiment, all solutions were bubbled with Carbogen (5% $CO_2$, 95%$O_2$). The VTA slices were recovered in a pre-warmed NMDG-HEPES aCSF solution at 34 °C and were subjected to a stepwise $Na^+$ reintroduction. Recovered slices were then transferred to a HEPES aCSF holding solution (in mM: NaCl, 92; KCl, 2.5; $NaH_2PO_4$, 1.2; $NaHCO_3$, 30; HEPES, 20; glucose, 25; sodium ascorbate, 5; thiourea, 2; sodium pyruvate, 3; $MgSO_4$, 2; $CaCl_2$, 2) at room temperature for at least 1 h prior to imaging. Imaging experiments were performed in recording aCSF (in mM: NaCl, 124; KCl, 2.5; $NaH_2PO_4$, 1.2; $NaHCO_3$, 24; HEPES, 5; glucose, 12.5; $MgSO_4$, 2; $CaCl_2$, 2).

*Imaging acute slices under bath perfusion of fentanyl*

Slices were transferred to an imaging chamber (Warner Instruments) and held in place using a harp (Warner Instruments) during bath perfusion at ~2 mL/min of aCSF bubbled with Carbogen (5% $CO_2$, 95%$O_2$) in gravity-fed syringes. The slices were imaged on an upright fluorescence microscope (Olympus BX50WI) equipped with a blue LED, filters, a dichroic mirror for GFP excitation/emission, and a CMOS camera (Hamamatsu ORCA-03G). First, the slice was visualized through a 4x (NA 0.10) objective to locate the VTA. Then fluorescence imaging was performed using a 40x (NA 0.80) water immersion objective and excitation from a blue LED. During the imaging experiment, the slice was bathed in aCSF, the solution was changed to 1 $\mu$M fentanyl, the drug was washed out, and then a pulse of aCSF from another syringe was applied as a pH control. No appreciable pH artifact was observed for the acute slice. Bath exchange time was ~10 s. No fluorescence was observed on the slice ipsilateral to the side of the injection site. Data were accepted from a slice that maintained its position throughout the imaging experiment, requiring no corrections.

*in vivo* iFentanylSnFR2.0 Recordings

*Surgical procedures for fiber photometry*

Pre-prepared optical fibers (diameter = 400 $\mu$m, NA 0.39) fixed in cannula were purchased from ThorLabs and cut to 4.9 mm using a ruby scribe. The fibers were inspected for their light transmittance and reflectance before use in surgery. Mice aged 8-12 weeks underwent anesthesia induction at 5% isoflurane mixed with oxygen and were maintained at 1-2% isoflurane during the surgery. Mice were kept on a heating pad during the surgery to maintain body temperature. The mice were mounted on a small animal stereotaxic apparatus (Stoelting) equipped with ear bars, and their skulls were leveled to place lambda and bregma on the same z-axis. A local anesthetic was applied, and the scalp was sterilized before beginning the surgery. The VTA was targeted for viral injection as described for the acute slice experiments using the same coordinates: AP -3.30; ML +0.40; DV −4.30. Viruses encoding functional and null versions of iFentanylSnFR2.0 were injected in different groups of animals. After the viral injection, the fiber optic cannula was mounted onto an adapter attached to the stereotactic rig, and the position of the fiber tip was calibrated with respect to lambda and bregma. The fiber was lowered into position 100 $\mu$m above the viral injection site over the course of ~5 min. The fiber was then secured to the skull with dental cement (Lang Dental). After the surgery, mice were placed in a new cage with a heating pad underneath half of the cage and observed until they regained movement. Thereafter, the mice were checked daily for a routine recovery. The mice were allowed 2 weeks of recovery before behavioral studies.

*Fiber photometry*

Bulk fluorescence signals from the VTA were measured using a fiber photometry setup as previously described (Augustine 2018). Briefly, collimated light from 490 nm and 405 nM light-emitting diodes (ThorLabs, M490F1 and M405F1) was directed via a tether to the optical cannula. The excitation light power at the end of the patch cord was measured to be < 75 μW for all experiments. The fluorescence output was focused onto a silicon femtowatt photoreceiver (Newport, Model 2151). A real-time processor (Tucker-Davis Technologies) performed modulation and demodulation using a MATLAB script previously reported by Augustine et al. (2018). This method was applied to all animals, including the negative control experiment using the null mutant, iFentanylSnFR2.0-436L.

*Videography rig*

A custom light and camera fixture was constructed for consistent videography and lighting during photometry experiments. A ceiling was constructed with two LED light bars (200 lumens, 4000K color, Defiant) fixed on each side of a 1080p camera (Logitech c920). The ceiling was fixed above an open arena (12” x 6” x 8”). A white paper towel was placed beneath the arena’s mesh floor, and a flat white background was placed behind two of the three transparent walls (the fourth wall is solid metal, and the other transparent wall was left unobstructed for the side view camera). A duplicate 1080p camera was positioned to capture the side view of the animal.

*Behavioral experiments involving photometry alongside videography*

Animals at 2-4 weeks post-surgery (10-12 weeks of age) were used for photometry plus videography experiments. All animals observed were opioid-naïve prior to the experiment. Animals were acclimated to the tether and the lit arena for 30 minutes before beginning the experiment. The photometry signal was verified, and the illumination power was set during this period. Each experiment consisted of a ~30 min baseline period, IP injection, and 3.5 h post-IP. Photometry and video recording were performed throughout the experiment uninterrupted, including the time spent handling and injecting the animal.

Machine vision applied to behavioral video

The recorded mouse behavior video was processed and analyzed using DeepLabCut (DLC) (Mathis et al., 2018) to extract the positions of the body parts of the animal in each frame. A separate DLC model was trained for each of the top and side views. Frames were extracted from videos using k-means clustering. The experimenter manually labeled the key points (annotated below). In the “top view”, the key points were the nose, head, fiber, ear_left, ear_right, neck, mid_1, mid_2, mid_3, tail_base, and tail_mid. Only the fiber and mid_1 key points were used for analyses from the top view. In the “side view”, the key points were the fiber, neck, body_mid, tail_base, tail_1-5, and tail_end. The tail_2-5 were placed as a series of bisections (i.e., tail_2 is halfway between tail_base and tail_end, tail_3 is halfway between tail_2 and tail_end, tail_4 is halfway between tail_3 and tail_end, and tail_5 is halfway between tail_4 and tail_end. A convolutional neural network, ResNet-50, was trained on the labeled frames augmented with imgaug. DLC was also used to generate movies of the input videos with the labeled points in frames where confidence exceeded ‘pcutoff’ = 0.7.

A single side camera allows for a quantitative measure of the Straub tail: the routine computes the tail angle when confidently visible. The tail may be hidden when the animal sits in a corner or obscured as the animal turns to face the camera. Mathis et al. have noted that DeepLabCut can be used to determine not only the location of a key point but also if a key point is not visible in a frame. The side view necessarily considers the tail against various backgrounds (white background, metal, and the animal itself), whereas the top view key points are inherently consistent. However, when animals display Straub tail, they are typically moving or, less often, stalled in a corner. In both cases, the tail is held against the white background, providing a clear view of all the key points frequently enough to capture the dynamics of the Straub tail.

*Histology*

Mice were euthanized with an intraperitoneal injection of 100 mg/kg of euthasol. Transcardial perfusion was performed with 1x PBS followed by 4% paraformaldehyde in 1x PBS. The brain was removed from the skull and

submerged in 4% PFA for 24 hours. The fixed brain was then rinsed three times with 1x PBS and sliced into 100 $\mu$m sections on a vibratome (Leica VT1200).
Fixed brain sections were incubated with 0.2 % Triton X-100 and 10% donkey serum in 1x PBS pH 7.4 on an orbital shaker at room temperature for 1 h. The slices were then incubated with primary anti-tyrosine hydroxylase antibody (Sigma-Aldrich AB1542) (1:200) in 10% goat serum with 0.2 % Triton X-100 overnight at 4 °C with gentle shaking. The slices were washed with three rinses with 1x PBS pH 7.4 for 5 min each and then incubated with donkey anti-sheep (IgG) secondary antibody conjugated to Alexa Fluor® 568 (Abcam, ab175712) (1:500) in 10% goat serum and 0.2 % Triton X-100 with gentle shaking at room temperature 6 h. Slices were then rinsed three times in 1x PBS pH 7.4 for 5 min each. Primary anti-GFP antibody (Abcam ab13970) (1:200) and secondary Goat Anti-Chicken IgY H&L Alexa Fluor® 488 (Abcam ab150169) (1:500) antibody were subsequently applied sequentially to stain the slices using the same procedure using donkey serum in place of goat serum. The stained slices were mounted onto glass microscope slides with coverslips using a mounting medium containing DAPI (Vector Laboratories),

*Fixed slice imaging*

The fixed slices were imaged using a Zeiss LSM980 confocal microscope to verify fiber placement above the VTA. A 2.5x objective was used to visualize entire slices and determine the slice with the deepest fiber tract to determine the position of the fiber tip. The region adjacent to the end of the fiber tract was imaged using a 20x (0.8 NA) objective and three channels in separate tracks, optimized by Zeiss Zen software:
AlexaFluor488, anti-GFP: 493 nm excitation, 508-578 nm emission, GaAsP-PMT detector
AlexaFluor568 settings, anti-Th: 577 nm excitation, 587-693 nm emission, GaAsP-PMT detector
DAPI: 353 nm excitation, 408-506 nm emission, Multialkali-PMT

Slice anatomy was checked against (Paxinos and Franklin) to verify the correct fiber position above the VTA. The Th-positive region provided a secondary validation of correct targeting. All reported animals showed a robust expression of the biosensor in the VTA and successful fiber positioning above the VTA.

#### Maze Foraging Paradigm

*Maze construction, lighting, and videography*

A labyrinth maze and lighting setup previously constructed and reported by Rosenberg et al. (2021) was used without modification in this work. The details of the apparatus from Rosenberg et al. are summarized here: the maze has six levels of T-junctions (each a left/right decision point) within a 24” x 24” layout. The passageways were 1.5” wide, and the floor-to-ceiling distance was 2”. The floor was constructed from IR-transparent acrylic. A water port (Sanworks) was installed at the peripheral end of one path and could be activated by nose-poking to break an IR beam. The port was calibrated to provide ~30 µL of drinking water for each nose-poke. A Bpod behavior box (Sanworks) was used to record all nose poke activity at the port and enforce a 90-second timeout period where subsequent nose pokes would not administer water. The port was also connected to an indicator IR LED that would flash for 1 second if water was administered. A video camera (c920, Logitech) was fixed below the maze to capture the animal’s movements within the maze and the indicator IR LED. Several IR illuminators (arrays of 12 IR LEDs) were placed throughout the room to provide an even lighting of the maze and contrast outlining the animal’s body. Three of these illuminators were placed underneath the maze, pointed at a 45-degree angle, to produce contrast between the maze floor and the animals’ footpads. The maze was slotted into a fixed frame, and the IR lights were fastened to arms on this frame so that the positions of all components were maintained across all experiments. A regular home cage was connected to the entry of the maze via a tube (3 cm in diameter, 1 meter long).

*Foraging experiments using the maze*

Wildtype C57Bl/6 (Charles River) mice were used for foraging experiments and were naïve to any manipulation or. Experiments were conducted for 10 h within the regular dark cycle of the mice. Mice were divided into two cohorts: the “intrinsic” cohort received food and water ad libitum before and during the experiment and were allowed to enter the maze with no water port. The “water-deprived” cohort was housed in a cage with the water pack removed 21 h prior to the experiment. All animals were singly housed for two days prior to the experiment. The water-deprived animals received an IP injection of either saline, 0.1 mg/kg fentanyl, or 1.0 mg/kg fentanyl and were placed in the home cage connected to the maze. The intrinsic cohort received an IP of either saline or

1.0 mg/kg fentanyl. Video of the maze was recorded continuously for 10 h. After the experiment, mice were returned to their regular home cage with food and water provided ad lib. The maze was cleaned with 70% EtOH in between animals and, once a week, was washed with a detergent.

##### Computational studies of proteins

###### *Docking*

All docking experiments were performed using AutoDock Vina (Eberhardt 2021 and Trott 2010) and repeated a method we previously used to dock an opioid into the crystal structure of iNicSnFR3a (PDB: 7S7T)) (Muthusamy 2022). The protein structure and ligands were prepared in AutoDockTools by removing water, adding polar hydrogens, and assigning the Gasteiger charges. The ligands were allowed torsional freedom in the docking routine. The docking figures and measurements between atoms were generated with PyMol (version 2.5.0, Schrodinger).

###### *Structure generation in AlphaFold2 and alignment*

The structures of the periplasmic binding protein portion of iFentanylSnFR2.0 was generated using AlphaFold2 executed through ColabFold (Merdita 2022; Jumper 2021). The predicted structure of iFentanylSnFR2.0's PBP and the crystal structure of iNicSnFR3a's PBP (PDB: 7S7W) were aligned in ChimeraX using the Needleman-Wunsch sequence alignment algorithm, and the BLOSUM95 substitution matrix was used to determine the RMSD between the two structures.

#### Quantification and Statistical Analysis

##### Dose response analysis for plate reader experiments

The biosensor's response was is expressed as ΔF/F0 and was calculated for each application of [ligand] where $\Delta F/F_0 = (F_{drug+biosensor} - F_{biosensor})/F_{biosensor}$ and each fluorescence readout (each 'F') was averaged across three replicates at a given [ligand]. The error is shown as the standard error of the mean where n = 3. The resulting $\Delta F/F_0$ vs. [ligand] set was fit using the Hill equation in Origin 9.2 (OriginLabs). Where the biosensor demonstrated a linear dose-response (i.e., [ligand] << $EC_{50}$), the data were fit with a linear regression with no y-axis restriction. This slope determined by the linear fit is termed the "S-Slope" and is expressed in the units of inverse concentration.

##### Principal component analyses (PCA)

This work performed two distinct PCAs. In both cases, the vectors were normalized before the PCA step using the "StadardScaler" function from scikit-learn's "preprocessing" package, and the PCA function was called from scikit-learns's "decomposition" package.

Figure 1H used RDKit to perform chemoinformatics and utilized PCA to display the groupings of chemical structures. The opioids used in this work were converted to their Morgan fingerprint vectors using RDKit's built-in functions. The groupings were drawn by the experimenter based on conventional opioid pharmacological classes, and the PCA results faithfully recapitulated the various classes shown in Figure 1H.

In Figure 2A, PCA was performed on the S-Slope heatmap result from Figure 1G to identify outliers in their binding mode against the cholinergic biosensor library.

##### Dose responses live cell experiments

This method applies to all live cell imaging time series data (culture and acute slice). Cells resolved in the focal plane along with a reference blank space were traced using the region of interest (ROI) tool in ImageJ. The "Time Series Analyzer" plugin determined the average pixel intensity in each ROI. These time series data were analyzed in Origin 9.1 (OriginLabs): the background ROI value was subtracted from each cell ROI to give F. A baseline was drawn from the initial and final periods, providing $F_0$. The difference in the response and the baseline gave ΔF and $\Delta F/F_0$ was plotted vs time. The steady-state response was determined by taking the average of final 40 frames (10 s) for each drug dose application. These steady-state responses were fit with either a linear regression or Hill fit.

##### Fiber photometry

ΔF/F was calculated using a MATLAB code described by the Deisseroth lab (Lerner et al. 2015). The data for the 405 nm and 490 nm channels were processed by applying a 1.8 Hz low-pass filter and scaling the 405 nm signal to the 490 nm signal using a linear function. ΔF/F was calculated as (raw 490 nm signal – fitted 405 nm signal) / (fitted 405 nm signal). The fluorescence data were subjected to a 1 min FFT filter for the plots.

##### Open arena machine vision outputs

A custom Python script was written to analyze the outputs from the analysis of the open arena videos processed by DeepLabCut. The script largely uses the scipy and statsmodels libraries for signal processing and statistical analyses and matplotlib for plotting. Each animal has three data inputs in this notebook: "top view" DLC output .csv, "side view" DLC output .csv, and the processed photometry signal .csv. First, these files are read in for each animal. Two additional inputs are read: a master spreadsheet listing timings for LED on/off and IP events for each animal and an image of the top view video to set reference coordinates. The LED on and off times set a global time reference when plotting photometry against behavior video results. Only frames where the labels of interest had a confidence score of > 0.6 were used for analysis. The "top view" outputs were used for all behavioral analyses except for the "Straub tail" angle measure, which is the only metric that uses the "side view" outputs.

The velocity (Fig 5C) and total distance traveled (Fig 6A) were calculated from the displacement of the "mid_1" body part in the "top view". The velocity was calculated for a rolling window of 100 frames (3.3 s). The resulting trace was Gaussian smoothed with σ = 1800 frames (60 s) for the plot in Fig 5C.

The circling (Fig 6B) and nose-in-corner (Fig 6C) metrics used the cage corners and midpoints in between the corners as landmarks. A circling bout was defined as the mid_1 body part passing within XXX cm from each reference point in sequence (i.e., tracing the cage's perimeter) within 30 s. A nose-in-corner bout was defined as the fiber label held within a threshold of 3 cm from a cage corner point for greater than 3 sec. Fig 6C and 6B plot histograms of the fraction of time spent in circling bouts and nose-in-corner bouts, respectively, for each 5 min bin.

The "Straub tail" was operationalized as the angle between the tail and the body. The body vector was defined as the vector from the "tail_base" label to the fiber label, and the tail vector was defined as the vector from the "tail_base" to "tail_4". The tail angle was set to NaN for frames where the tail was hidden, or labels did not have sufficient confidence. The resulting trace was Gaussian smoothed with σ = 1800 frames (60 s) for the plot in Fig 5C.

##### Maze foraging behavior

The analysis of the maze experiments replicated those previously reported by Rosenberg et al. 2021. Briefly, the raw video was analyzed in grayscale to determine the animal's trajectory (key points: nose, feet, tail base, and mid-body). This analysis provided the x-y coordinates vs. time used in the previously reported notebooks (https://github.com/markusmeister/Rosenberg-2021-Repository). This work had up to three experimental groups, so, where appropriate, the code for the plots were adapted to include a third group.

#### Methods References:

Augustine, Vineet, Sertan Kutal Gokce, Sangjun Lee, Bo Wang, Thomas J. Davidson, Frank Reimann, Fiona Gribble, Karl Deisseroth, Carlos Lois, and Yuki Oka. "Hierarchical neural architecture underlying thirst regulation." *Nature* 555, no. 7695 (2018): 204-209.

Jumper, John, Richard Evans, Alexander Pritzel, Tim Green, Michael Figurnov, Olaf Ronneberger, Kathryn Tunyasuvunakool et al. "Highly accurate protein structure prediction with AlphaFold." *Nature* 596, no. 7873 (2021): 583-589.

Challis, Rosemary C., Sripriya Ravindra Kumar, Ken Y. Chan, Collin Challis, Keith Beadle, Min J. Jang, Hyun Min Kim et al. "Systemic AAV vectors for widespread and targeted gene delivery in rodents." *Nature protocols* 14, no. 2 (2019): 379-414.

Eberhardt, J., Santos-Martins, D., Tillack, A.F., Forli, S. (2021). AutoDock Vina 1.2.0: New Docking Methods, Expanded Force Field, and Python Bindings. Journal of Chemical Information and Modeling.

Paxinos, George, and Keith BJ Franklin. *Paxinos and Franklin's the mouse brain in stereotaxic coordinates*. Academic press, 2019.

Kille, Sabrina, Carlos G. Acevedo-Rocha, Loreto P. Parra, Zhi-Gang Zhang, Diederik J. Opperman, Manfred T. Reetz, and Juan Pablo Acevedo. "Reducing codon redundancy and screening effort of combinatorial protein libraries created by saturation mutagenesis." *ACS synthetic biology* 2, no. 2 (2013): 83-92.

Mirdita, Milot, Konstantin Schütze, Yoshitaka Moriwaki, Lim Heo, Sergey Ovchinnikov, and Martin Steinegger. "ColabFold: making protein folding accessible to all." *Nature methods* 19, no. 6 (2022): 679-682.

Muthusamy, Anand K., Charlene H. Kim, Scott C. Virgil, Hailey J. Knox, Jonathan S. Marvin, Aaron L. Nichols, Bruce N. Cohen, Dennis A. Dougherty, Loren L. Looger, and Henry A. Lester. "Three mutations convert the selectivity of a protein sensor from nicotinic agonists to S-methadone for use in cells, organelles, and biofluids." *Journal of the American Chemical Society* 144, no. 19 (2022): 8480-8486.

Trott, O., & Olson, A. J. (2010). AutoDock Vina: improving the speed and accuracy of docking with a new scoring function, efficient optimization, and multithreading. Journal of computational chemistry, 31(2), 455-461.

Rosenberg, Matthew, Tony Zhang, Pietro Perona, and Markus Meister. "Mice in a labyrinth show rapid learning, sudden insight, and efficient exploration." *Elife* 10 (2021): e66175.

Schindelin, J., Arganda-Carreras, I., Frise, E., Kaynig, V., Longair, M., Pietzsch, T., … Cardona, A. (2012). Fiji: an open-source platform for biological-image analysis. Nature Methods, 9(7), 676–682. doi:10.1038/nmeth.2019

Ting, Jonathan T., Brian R. Lee, Peter Chong, Gilberto Soler-Llavina, Charles Cobbs, Christof Koch, Hongkui Zeng, and Ed Lein. "Preparation of acute brain slices using an optimized N-methyl-D-glucamine protective recovery method." *JoVE (Journal of Visualized Experiments)* 132 (2018): e53825.

**Biosensor amino acid sequences:**

His$_6$ tag
HA tag
PBP lobes
Linkers
cpGFP
Myc tag

**iFentanylSnFR2.0:**

HHHHHHGYPYDVPDYAGAQPARSANDTVVVGSANFTEGIIVANMVAEMIEAHTDLKVVRKLNLG
GGNVNFEAIKRGGANNGIDIYVEYTGHGLVDILGFPEPNVYITADKQKNGIKANFKIRHNVEDG
SVQLADHYQQNTPIGDGPVLLPDNHYLSTQSVLSKDPNEKRDHMVLLEFVTAAGITLGMDELYK
GGTGGSMSKGEELFTGVVPILVELDGDVNGHKFSVRGEGEGDATNGKLTLKFICTTGKLPVPWP
TLVTTLTYGVQCFSRYPDHMKQHDFFKSAMPEGYVQERTISFKDDGTYKTRAEVKFEGDTLVNR
IELKGIDFKEDGNILGHKLEYNFPPPISTDPEGAYETVKKEYKRKWNIVWLKPLGFNNTGTLTV
KDELAKQYNLKTFSDLAKISDKLILGATMFFLEAPDGYPGLQKLYNFKFKHTKSMDMGIRYTAI
DNNEVQVIDAWATDGLLVSHKLKILEDDKAFFPPYYAAPIIRQDVLDKHPELKDVLNKLANQIS
GEEMQKLNYKVDGEGQDPAKVAKEFLKEKGLILQVDEQKLISEEDLN

**Table XXX**: Substitution mutations in exemplary opioid biosensors with respect to [insert parent sequence reference]

| **Biosensor** | **Opioid ligand** | **Mutations** |
|---|---|---|
| iFentanylSnFR1.0 | Fentanyl | K10A, Q15G, T43E, T68H, T325S, K330G, D341R, Y357G, A360T, E391F, R395A, V405L, F436W, H455A, D490L |
| iFentanylSnFR2.0 | Fentanyl | K10A, Q15G, T43G, T68H, T325S, K330G, D341R, Y357G, A360T, E391F, R395A, V405L, F436W, H455A, D490G |
| iTapentadolSnFR1.0 | Tapentadol | K10I, N11E, Q15G, T43E, T68H, T325S, K330G, D341R, A360T, E391F, R395G, V405L, F436A, H455A, D490L |
| iS-methadoneSnFR1.0 | S-methadone | K10I, N11V, Q15G, T43E, T68H, T325S, K330G, D341R, A360T, E391F, R395G, V405L, H455A, D490G |
| iLevorphanolSnFR1.0 | Levorphanol | K10I, N11P, Q15G, T43V, T68H, K330G, D341R, A360T, E391F, R395G, V405L, F436W, H455W, D490L |
| iBRL52537SnFR lead 1 | BRL52537 | K10I, N11E, Q15G, T43E, T325S, K330G, D341R, A360T, E391F, R395G, V405L, F436W, H455A, D490L |
| iBRL52537SnFR lead 2 | BRL52537 | K10I, N11E, Q15G, T43E, T325S, K330G, D341R, A360T, E391F, R395G, V405L, F436W, H455A, D490L |
| iBRL52537SnFR lead 3 | BRL52537 | K10I, N11E, Q15G, T43E, T325S, K330G, D341R, A360T, E391F, V405L, F436W, H455A, D490L |
| iTramadolSnFR lead 1 | Tramadol (racemic) | K10I, N11E, Q15G, T43E, T68H, T325S, K330G, D341R, A360T, E391F, R395G, V405L, F436T, H455A, D490L |

| | | |
|---|---|---|
| iTramadolSnFR lead 2 | Tramadol (racemic) | K10I, N11E, Q15G, T43E, T68H, T325S, K330G, D341R, A360T, E391F, R395G, V405L, F436C, H455A, D490L |
| iButorphanolSnFR lead 1 | Butorphanol | K10I, N11E, Q15G, T43E, T325S, K330G, D341R, A360T, E391F, R395G, V405L, F436W, H455A, D490L |
| iNorfentanylSnFR lead 1 | Norfentanyl | K10I, N11E, Q15G, T43E, T325S, K330G, D341R, A360T, E391F, V405L, F436W, H455A, D490L |
| iSufentanilSnFR lead 1 | Sufentanil | K10I, Q15G, T43E, T68H, T325S, K330G, D341R, Y357G, A360T, E391F, R395M, V405L, F436W, H455A, D490L |
| iMorphineSnFR lead 1 | Morphine | K10I, N11P, Q15G, T43I, T68H, K330G, D341R, A360T, E391F, R395G, V405L, F436W, H455G, D490L |
| iMorphineSnFR lead 2 | Morphine | K10I, N11P, Q15G, T43I, T68H, K330G, D341R, A360T, E391F, R395G, V405L, F436W, H455Q, D490L |
| iCodeineSnFR lead 1 | Codeine | K10I, N11F, Q15G, T43V, T68H, K330G, D341R, A360T, E391F, R395G, V405L, F436W, H455A, D490L |
| iCodeineSnFR lead 2 | Codeine | K10I, N11L, Q15G, T43V, T68H, K330G, D341R, A360T, E391F, R395G, V405L, F436W, H455A, D490L |
| iCodeineSnFR lead 3 | Codeine | K10I, N11P, Q15G, T43V, T68H, K330G, D341R, A360T, E391F, R395G, V405L, F436W, H455A, D490L |
| iHydromorphoneSnFR/iHydrocodone lead 1 | Hydrocodone and Hydromorphone | K10I, N11D, Q15G, T43V, T68H, K330G, D341R, A360T, E391F, R395G, V405L, F436W, H455A, D490L |
| iNaltrexoneSnFR lead 1 | Naltrexone | K10V, N11E, Q15G, T43E, T325S, K330G, D341R, A360T, E391F, R395G, V405L, F436W, H455A, D490L |
| iNaltrexoneSnFR lead 2 | Naltrexone | K10A, N11E, Q15G, T43E, T325S, K330G, D341R, A360T, E391F, R395G, V405L, F436W, H455A, D490L |
| iNaltrexoneSnFR lead 3 | Naltrexone | K10M, N11E, Q15G, T43E, T325S, K330G, D341R, A360T, E391F, R395G, V405L, F436W, H455A, D490L |
